# HTRA3 protease-chaperone stabilizes cathepsin B for mitochondrial POLG1 depletion in human cell ageing

**DOI:** 10.1101/2025.07.21.647696

**Authors:** Cristina Fernández Molina, Laurent Chatre, Benjamin Montagne, Audrey Salles, Alain Sarasin, Clément Crochemore, Miria Ricchetti

## Abstract

The maintenance of mitochondrial proteins homeostasis, which is essential for proper cell function, is affected in pathophysiological ageing, yet several underlying mechanisms remain unexplored. We show that in normal and accelerated ageing cells, POLG1, the enzyme responsible for mitochondrial DNA replication, is degraded by the protease cathepsin B, which is overexpressed, escapes from lysosomes, and is stabilized by the chaperone activity of another protease, HTRA3. This degradation is in part counteracted by RAC1, a small GTPase also stabilized by HTRA3. POLG1 depletion, that occurs in progeroid Cockayne syndrome and senescent cells, is linked respectively to impairment or downregulation of the CSB protein, which promote cellular senescence. Our experiments in engineered cells, demonstrate that senescence itself, and not the absence of CSB, triggers the accumulation of cathepsin B and HTRA3, leading to POLG1 degradation. In summary, we uncover a complex, multi-step process that controls the degradation of POLG1 in mitochondria, a process that is activated by cell senescence and becomes more pronounced in Cockayne syndrome cells, providing new insight in the regulation of mitochondrial proteostasis in ageing and progeroid disorders.

## Introduction

POLG1 is the catalytic subunit of DNA polymerase γ (Polγ), responsible for the replication and maintenance of mitochondrial DNA (mtDNA) in animal cells (1). The mechanisms governing POLG1 degradation, which is enhanced in progeroid Cockayne syndrome cells (2) and senescent cells (3), remain uneplored.

Polγ functions as a heterodimer composed of POLG1, associated with a dimeric accessory subunit, POLG2. POLG1 has three catalytic activities: 5′−3′ DNA polymerase (DNA synthesis), 3′−5′ exonuclease (proof-reading), and 5′-deoxyribose phosphate lyase activity, which is required for base excision repair, a DNA repair process that is active also in mitochondria. POLG1-defective activity can cause mtDNA depletion and accumulation of point mutations and deletions in the mitochondrial genome.

While mutations in POLG1 are linked to a variety of mitochondrial diseases that range from prenatally-fatal conditions and severe infantile onset disorders to late-onset, milder conditions (4), its role in ageing is increasingly evident. Alteration in POLG1 proof-reading and polymerase activity are associated with premature aging (5, 6), and a significant age-related decline in the mtDNA content is linked to physiological decline (7-10). Low levels of POLG1 have been associated with pathophysiological ageing conditions (2, 3), yet the mechanisms underlying the degradation of POLG1, particularly in the context of aging, remain largely unexplored,

We previously reported reduced levels of POLG1 in cells from patients with Cockayne syndrome (CS), a rare autosomal recessive disorder characterized by severe precocious ageing, neurodegeneration and, in the majority of cases, cutaneous photosensitivity (11). CS results from mutations in either *CSB* or *CSA*, coding for CSB and CSA, respectively, which are involved in the transcription-coupled nucleotide excision repair (TC-NER) pathway that repairs UV-induced DNA damage, and also act as transcription factors, and CSB is a chromatin remodelling factor (12). We showed that POLG1 depletion, leading to reduced OXPHOS in CS cells, was mediated by overexpression of the HTRA3 protease, and triggered by oxidative and nitrosative stress (11). This process was not observed in cells derived from patients with the related UVSS syndrome, which can also be due to CSA or CSB impairment, but is not characterized by precocious ageing, despite DNA repair defects and photosensitivity (13). These data suggest that POLG1 degradation is linked to the progeroid defect, and is independent of DNA repair impairment. POLG1 depletion leads to reduced OXPHOS in CS cells (11). Several mitochondrial defects have been associated with CS (11, 14), which shares characteristics of mitochondrial diseases like a progressive nature, and a large spectrum of onsets and clinical severity, linking the mitochondrial defects to the precocious ageing phenotype.

We also demonstrated that CS alterations, *i.e.* HTRA3 overexpression/POLG1 depletion/mitochondrial dysfunction, are recapitulated during replicative senescence of normal cells (3). Replicative senescence, characterized by a proliferative arrest of metabolically active cells, is a hallmark of ageing (15), marked by the accumulation of senescent cells, which contribute to tissue dynfunction and age-related diseases by withdrawing stem cells from the regeneration process (16). POLG1 degradation during senescence mirrors the mitochondrial dysfunction observed in CS and may represent a broader mechanism by which mitochondrial decline contributes to aging (14).Replicative senescence is normally induced by DNA damage generated during DNA replication, or persistent DNA damage response (DDR) resulting from telomere shortening, and leading to the stabilization of the transcription factor p53 and expression of the cyclin-dependent kinase inhibitor p21(17). We showed that in normal cells replicative senescence, and thereby POLG1 depletion, is triggered by CSB depletion that exposes the *p21* promoter and allows the expression of p21, in a p53-independent manner (3).

In this study, we identify cathepsin B (CTSB) as the protease that degrades POLG1 upon stabilization by HTRA3. Despite its primary lysosomal role, CTSB accumulates in CS cells as well as senescent cells and localizes in mitochondria upon lysosome destabilization, directly impacting mitochondrial function. We also show that in normal cells CTSB-dependent POLG1 degradation depends on replicative senescence rather than CSB depletion, provideing a new patwhay for the regulation mitochondrial protein turnover in age-related conditions.

## Results

### CTSB interacts with POLG1 and HTRA3 in WT and CS cells and accumulates in CS cells

To assess the mechanism of HTRA3-dependent POLG1 depletion in CS, explorative Mass Spectrometry (MS) experiments of HTRA3 and POLG1 interactors suggested no direct HTRA3 and POLG1 interaction, and rather interaction of cathepsin B with POLG1 and, in most tested CS cells, also with HTRA3 (not shown). Catepsin B (CTSB) is a member of the family of lysosomal cysteine proteases. We thus hypothesized a POLG1 degradation process resulting from the action of two proteins, namely HTRA3 (previously demonstrated (2)), and CTSB (see scheme in Fig. 1A). To explore this possibility, we assessed the interactions of CTSB with HTRA3 by co-immunoprecipitation experiments in CSB-deficient (CS1AN-SV), and wild type (WT) (MRC5-SV) immortalized fibroblasts. CTSB is synthesized as an inactive precursor that is activated when it reaches the endosomal/lysosomal compartments(18). We observed that the active form of CTSB (of 25 kDa) co-immunoprecipitates with endogenous HTRA3 in CS fibroblasts, as predicted in our model, and also in WT cells (Fig. 1B). Co-immunoprecipitation experiments also showed interaction of CTSB with POLG1 (Fig. 1B). These experiments suggested upregulation of the CTSB protein in CS1AN-SV compared to MRC5-SV cells (Fig. 1B, Input), which was confirmed by RT-qPCR (*SI Appendix,* Fig. S1A), and immunofluorescence (IF) (*SI Appendix,* Fig. S1B and C) experiments. To assess whether accumulation of CTSB in CS cells actually translated into an increased proteolytic activity, we incubated cells with the Magic Red CTSB substrate that generates a cresyl violet fluorophore proportional to CTSB activity. We observed a significant increase in the protease activity in CS1AN-SV cells compared to MRC5-SV controls (*SI Appendix,*Fig. S1D and E).

**Figure 1.**
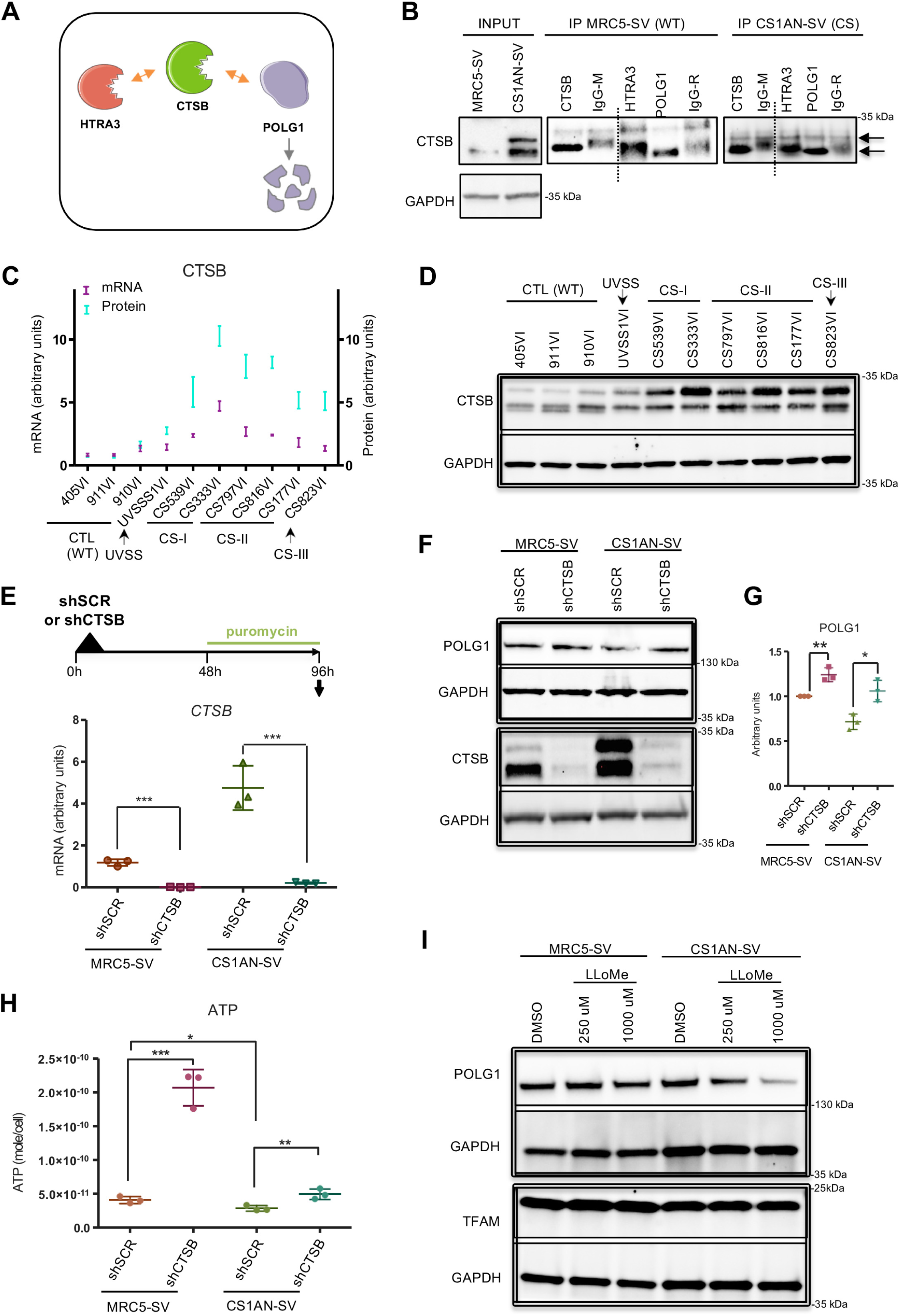
CTSB accumulates in CS cells and interacts with POLG1 and HTRA3 in WT and CS cells. (**A**) Schematic representation of protein interactions associated with POLG1 degradation. Orange arrows show interactions suggested by mass spectrometry analysis of HTRA3 and POLG1 interacting partners (not shown). (**B**) Co-immunoprecipitation assays showing interaction between CTSB and HTRA3 as well as CTSB and POLG1 in WT (MRC5-SV) and CS (CS1AN-SV) cell lines. Immunoprecipitation with Ig isotypes of the corresponding antibodies served as control. IP: immunoprecipitation. The 31-kDa CTSB band corresponds to the single-chain active CTSB form, and the 25 kDa to the heavy chain of the double-chain active CTSB form. CTSB single and double-chain active forms have been reported to have the same activity (52, 53). (**C**) mRNA levels of *CTSB* in primary WT (CTL), UVSS, and CS patient-derived fibroblasts compared to protein levels (SD, quantification of band intensity normalised to the respective GAPDH levels in three independent experiments, from *SI Appendix,* Fig. S1G). (**D**) WB of CTSB in primary WT, UVSS, and CS patient-derived fibroblasts. GAPDH (36 KDa) was used as a loading control for the WB. (**E**) Scheme of the CTSB silencing experiment (*upper panel*). RT-qPCR analysis of *CTSB* mRNA levels in MRC5-SV and CS1AN-SV cells expressing scramble control (shSCR) or *CTSB* shRNAs (*lower panel*). (**F**) Immunoblot of CTSB and POLG1 upon expression of shSCR or CTSB shRNAs, and the respective GAPDH as a control. (**G**) Quantification of POLG1 protein levels in n=3 independent experiments normalized with GAPDH. (**H**) OXPHOS (oligomycin-sensitive) activity in ATP synthesis per 10,000 cells. (**i**) WB analysis of POLG1 and TFAM (each with the loading control GAPDH) after 3 h treatment with LLoMe (250 µM or 1000 µM) followed by 1 h of drug withdrawal. RT-qPCR experiments: n=3; mean ± SD. \**p* ≤ 0.05, \*\**p* ≤ 0.01, \*\*\**p* ≤ 0.001; based on one-way ANOVA with post-hoc Tukey’s test *vs*. the mean of the controls 405VI, 911VI, 910VI (panel C) or the unpaired t test *vs.* shSCR (panels F, H).

These results prompted us to assess whether CTSB levels are high in CS patient-derived primary skin fibroblasts. In these cells we previously reported POLG1 depletion, whereas UVSS cells that are mutated but the carrier patient had not the progeroid phenotype, displayed normal levels of POLG1 as WT cells(11). CS has been classified in different forms with decreasing onset and clinical severity: CS-II, CS-I, CS-III (19). In cells derived from patients with the severe forms (CS-I and CS-II), transcript levels of *CTSB* were 1.8- to 4.7-fold higher than WT cells (Fig. 1C, purple, patient-derived cells are described in *SI Appendix,* Fig. S1F). CTSB expression was not higher than control in cells from the milder CS-III form (late-onset form of CS, allowing longer-term survival), and UVSS (not progeroid). CTSB protein levels assessed by Western blots (WB) were 5- to 10-fold higher in all CS patient-derived cells, including CS-III cells (Fig. 1D, *SI Appendix,* Fig. S1G), compared to WT or UVSS cells, confirmed by IF in selected representative cells (*SI Appendix,* Fig. S1H and S1I). Interestingly, the largest differences in CTSB content between WT and CS cells appeared at the protein rather than mRNA level, as highlighted by combining both sets of data (Fig. 1C), suggesting a stabilization of the CTSB protein or its mRNA in CS cells. To note, the expression of relevant mitochondrial proteases (*LONP1*, *CLPP*, *AGF3L2*, and *SPG7*) remained mostly unchanged in CS compared to control and UVSS1 cells (*SI Appendix,* Fig. S2 A-D).

Taken together, these results suggest interaction among HTRA3, CTSB, and POLG1 in WT cells, and at a larger scale in CS cells where CTSB accumulates. Accumulation of HTRA3 in CS cells was previously demonstrated (11).

### CTSB induces POLG1 protein depletion in WT and to a larger extent in CS cells

To verify whether CTSB is responsible for POLG1 depletion, we silenced *CTSB*. For that, MRC5-SV and CS1AN-SV cell lines were transduced with either a non-targeting shRNA (shRNA-scramble (SCR)) or a shRNA directed against *CTSB* (shCTSB). At day 4 following transduction, we observed more than 95% silencing of *CTSB* transcripts in both cell lines compared to the respective shSCR (Fig. 1E). Western blot analysis confirmed almost complete depletion of the CTSB protein (Fig. 1F). As predicted, CTSB silencing resulted in higher levels of POLG1 in MRC5-SV, and to a large extent in CS1AN-SV cells where the originally low levels of this polymerase returned as in control cells (Fig. 1F and G). Increase in POLG1 protein levels upon *CTSB* knockdown was further confirmed by immunofluorescence in WT and CSB-deficient cells (*SI Appendix,* Fig. S3A and B).

To assess the impact of increased/restored POLG1 levels on the mitochondrial function, we measured cellular ATP produced by mitochondrial oxidative phosphorylation (OXPHOS), using a luminescent assay in the presence and in the absence of oligomycin that blocks OXPHOS. As expected according to our working model, WT and CSB-deficient cells displayed an increase in mitochondrial ATP production upon CTSB knockdown, compared to the respective shSCR (Fig. 1H), with CS cells restoring the levels of (WT) controls. To note, in control MRC5-SV cells that originally carried physiological levels of POLG1, CTSB silencing resulted in a particularly large (about 5-fold) increase of mitochondrial ATP.

It has been reported that CS cells accumulate lysosomes with ruptured membranes(20), leading us to suggest that in these cells mature active CTSB leaks from lysosomes where this enzyme is mainly localized. To validate this hypothesis, we destabilized lysosomal membranes of control MRC5-SV (WT) cells with 250 µM or 1000 µM of L-leucyl-L-leucine methyl ester (LLOMe) for 3 hours, and indeed observed a dose-dependent decrease of the POLG1 protein content (Fig. 1I). The LLOMe treatment worsened POLG1 depletion in CS1AN-SV cells. POLG1 depletion upon LLOMe treatment seems to be specific to this protein, since the level of TFAM, another mitochondrial matrix protein implicated in mtDNA transcription and replication, remained unchanged under the same treatment (Fig. 1I). Taken together, these experiments show that CTSB is directly associated with POLG1 degradation in WT and, to a larger extent, in CS cells where this protease is present in larger amounts, and the process is enhanced upon destabilization of lysosomal membranes.

### CTSB is stabilized by HTRA3

Some members of the HtrA family, such as HTRA2, have been described to possess chaperone properties besides the protease activity (21). Since the chaperone-like activity of HTRA3 has been recently reported (22), we hypothesized that HTRA3, which is expressed at high levels in CSB-deficient cells, acts as a stabilizer of CTSB. Consistent with this notion, all tested CS patient-derived cells display high levels of CTSB (protein and mRNA, see above Fig. 1C and D), as well as high levels of HTRA3 (except AS177) as previously reported (11). CS823VI CS-III cells, that were not previously tested, also overexpress the CTSB protein (see above, Figs. 1C and D,Fig S1G), and show high levels of the HTRA3 protein (*SI Appendix,* Fig. S4A). To examine whether HTRA3 stabilizes CTSB, we performed RNA-mediated depletion of HTRA3, using the silencing strategy described above. *HTRA3* downregulation was effective in WT as well as CS cells (transcript knock-down of 72% in MRC5-SV and 56% in CS1AN-SV; Fig. 2A). Importantly, upon downregulation, *HTRA3* levels in CS cells were almost as low as in WT cells, which was confirmed at the protein level (Fig. 2B). In CS1AN-SV cells, HTRA3 knock-down reduced the abundant CTSB protein as early as 48h after shRNA transduction (Fig. 2B). Despite low native levels of the HTRA3 and CTSB proteins in WT cells, HTRA3 depletion led to a further (slight) decline in CTSB levels, consistent with CTSB and HTRA3 interacting not only in CS but also in normal cells.

**Figure 2.**
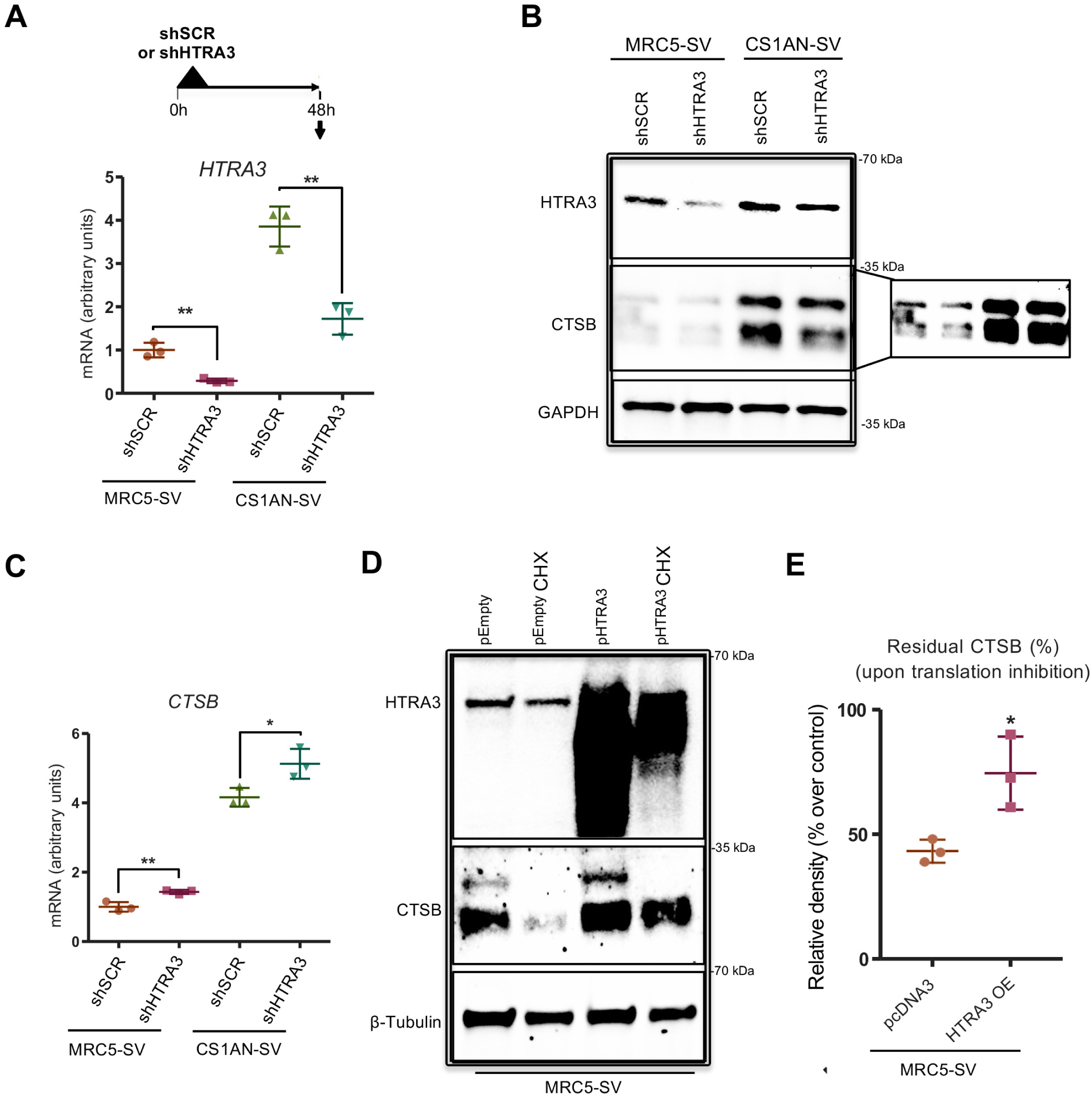
CTSB is stabilized by HTRA3. (**A**) Experimental set-up of the HTRA3 silencing experiment (*upper panel*). RT-qPCR of *HTRA3* in MRC5-SV and CS1AN-SV cells knocked-down for HTRA3 (shHTRA3) with respective scramble (shSCR) at 48 h after sh-RNA transduction (*lower panel*). (**B**) WB of HTRA3, CTSB and GAPDH (loading control) in the same cells as in panel a. A more exposed membrane for CTSB is shown on the right. (**C**) RT-qPCR of *CTSB* and after HTRA3 knock-down also at 48 h after sh-RNA transduction (**D**) WB of HTRA3 and CTSB in MRC5-SV cells transfected with a vector coding for HTRA3 (pHTRA3) or an empty vector (pEmpty), and treated or not with 30 µg/ml of cycloheximide (CHX) for 24 h. (**E**) The percentages of CTSB remaining after CHX treatment were determined by quantifying band intensities normalized to β-tubulin levels (loading control), and calculating the difference between the treated and the non-treated condition for both pEmpty vector and pHTRA3. RT-qPCR: n=3 independent experiments; mean ± SD. \**p* ≤ 0.05, \*\**p* ≤ 0.01, \*\*\**p* ≤ 0.001; based on unpaired t test comparisons *vs*. respective shSCR.

*HTRA3* knockdown did not lead to a decrease in *CTSB* transcripts, indicating that HTRA3 regulates CTSB post-transcriptionally, as we predicted. Rather, *HTRA3* knockdown increased *CTSB* transcripts (Fig. 2C), suggesting a homeostatic mechanism that compensates for decreased protein levels. To note, in control MRC5-SV cells, *HTRA3* silencing was maintained 96 h post-transduction, and consequently the levels of the CTSB protein remained low at this late time point (*SI Appendix,* Fig. S4B). Conversely, in CS1AN-SV cells, despite *HTRA3* RNA silencing (*SI Appendix,* Fig. S4C) the protein content increased back to normal (*SI Appendix,* Fig. S4B), suggesting stabilization of the HTRA3 protein in the long term in these cells. This event was accompanied by an increase in CTSB protein and mRNA content (*SI Appendix,*Fig. S4B and D), further supporting the notion of a direct correlation between HTRA3 and CTSB protein levels, and thereby CTSB stabilization by HTRA3.

In support of this model, ectopic overexpression of HTRA3 in MRC5-SV cells using a pcDNA3.1-based vector led to an increase of CTSB protein levels (Fig. 2D). Treatment of HTRA3 overexpressing cells for 24h with cycloheximide (CHX), a translation inhibitor, showed a decline in CTSB levels (Fig. 2D), as expected due to the blockage of protein synthesis. Interestingly, the fraction of residual CTSB was higher in HTRA3 overexpressing cells than cells transfected with the empty vector (Fig. 2E). This experiment indicates that, in the absence of new protein synthesis, HTRA3 overexpression maintains, at least a fraction of, the levels of CTSB, *i.e.,* reducing the degradation of CTSB, compatibly with the stabilization of this protein.

In conclusion, HTRA3 ablation reduced the CTSB protein content without affecting mRNA levels and, consistently, HTRA3 overexpression correlated with high CTSB protein levels and increased post-translational stability of this protein. Thus, these data reveal a direct correlation between HTRA3 levels and CTSB stability.

### Active CTSB is present in mitochondria

As CTSB is directly associated with POLG1 depletion, we investigated the subcellular localization of this interaction. POLG1 is a nuclear-encoded protein, translated in the cytosol, and imported into mitochondria. Western blots on the cytosolic and mitochondrial fractions after subcellular fractionation of MRC5-SV and CS1AN-SV cells showed that POLG1 was prevalently located in the mitochondrial fraction in both cell lines (Fig. 3A), as expected. CTSB appeared also prevalently located in the mitochondrial fraction, and in particular in CS1AN-SV cells where this protein is abundant. Thus, we reasoned that POLG1 degradation may take place directly in the organelle. In agreement with this notion, lower levels of POLG1 in whole-cell lysate in CS1AN-SV compared to MRC5-SV correlate with depletion of the protein in the mitochondrial, but not the cytosolic fraction (Fig. 3A). The other member of the putative HTRA3/CTSB/POLG1 complex, namely HTRA3, was also detected in the mitochondrial fraction (although prevalently in the cytosol), (Fig. 3B). This result is consistent with the localization of HTRA3 in the cytosol and mitochondria, as we previously reported with IF super-resolution microscopy analysis (3).

**Figure 3.**
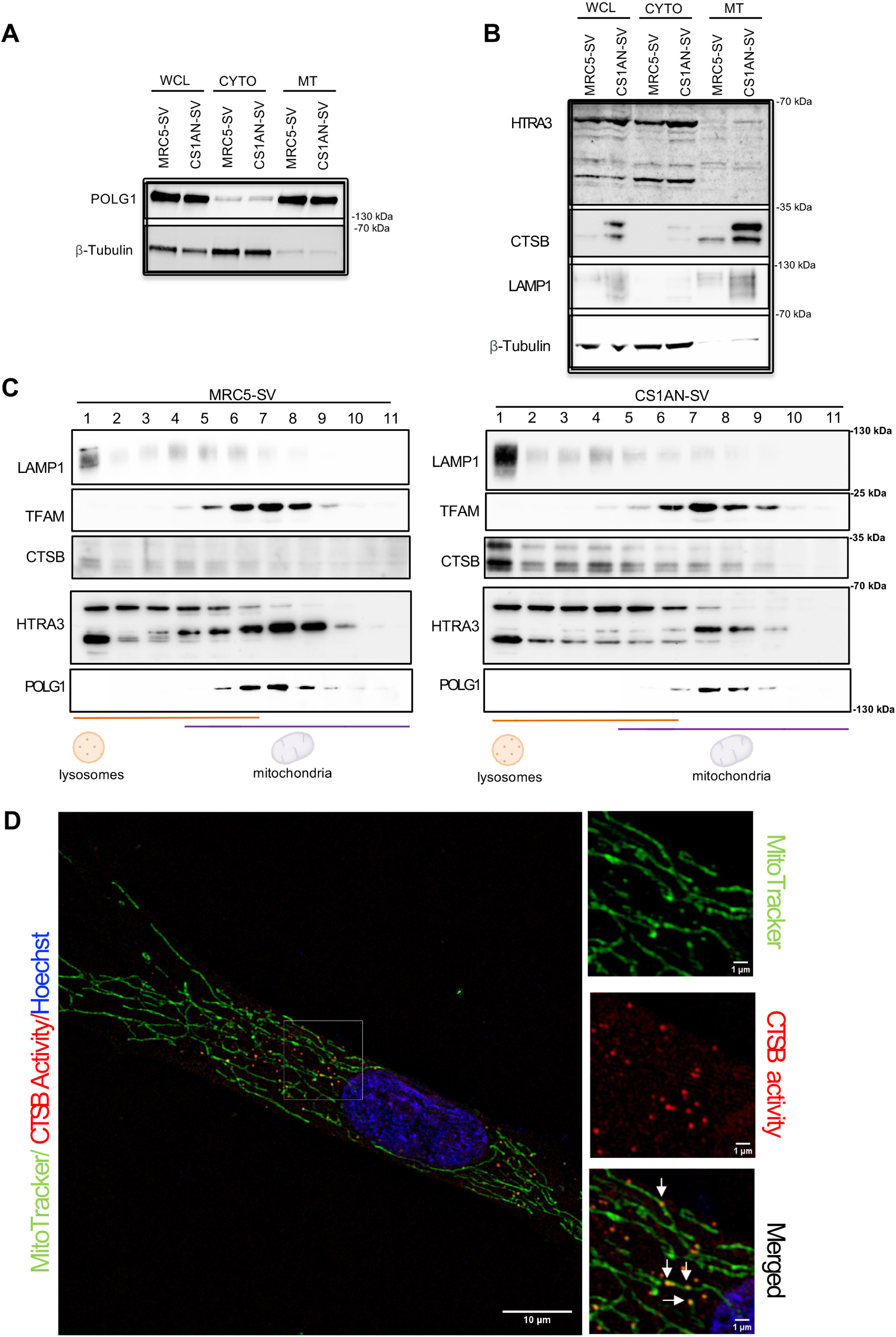
Active CTSB is present and in mitochondria. WB of POLG1 (**A**), and HTRA3, CTSB, and LAMP1 (**B**) in whole cell lysate (WCL), subcellular mitochondrial (MITO) and cytosolic (CYTO) fractions in MRC5-SV and CS1AN-SV cells. LAMP1 lysosomal marker, β-tubulin cytosolic marker, POLG1 mitochondrial marker. During stress, HTRA3 undergoes autocleavage that generates shorter forms (54), which are visible in panel B. Shorter HTRA3 isoforms were detected in our previous study in IMR90 fibroblasts but not, or were very faint, in CS cells (11); these isoform have not been detected in other WBs in the present study. The increased detection of the HTRA3 cleaved forms in fractionation experiments is possibly due to the longer time required for this experiment, with consequent HTRA3 auto-cleaving during manipulations. (**C**) Immunoblot analysis of LAMP1, TFAM, CTSB, HTRA3, and POLG1 in MRC5-SV and CS1AN-SV samples fractionated on an Optiprep density gradient. A clear separation is observed for lysosomes, as shown by the lysosomal marker LAMP1, mainly found in fraction 1, and mitochondrial markers, TFAM and POLG1, mainly founds in fractions 6–11. (**D**) Sublocalization of CTSB activity and mitochondria in CS1AN-SV with SIM (one plan of a z-stack acquisition for each cell). Mitochondria were visualized using MitoTracker (green), CTSB activity using Magic Red CTSB (red), and nuclei were stained with Hoechst (blue). Scale bars = 10 µm. The 1.5x magnification of a specific region is shown on the right with immunostaining for MitoTracker, CTSB activity, and merge (representative arrow for colocalization of CTSB activity/MitoTracker).

These experiments suggested that the POLG1/CTSB interaction, leading to degradation of the polymerase, takes place in mitochondria. However, we wondered whether the fractionation method efficiently separated lysosomes from mitochondria. Labelling of LAMP1, a lysosomal membrane protein, revealed that the enriched mitochondrial fraction actually contained lysosome components, and at a larger extent in CS1AN-SV cells (Fig. 3B). To corroborate these findings, we used OptiPrep density ultracentrifugation that accurately separates lysosomes from mitochondria (23) (Fig. 3C). Lysosomes were mainly recovered in fraction 1, as shown by the lysosomal marker LAMP1. POLG1 was detected in fractions 5 to 9, *i.e.,* in the mitochondrial fraction, as indicated by the position of the mitochondrial transcription factor TFAM that is located in the organelle matrix. HTRA3 was located in the lysosomal, mitochondrial, and intermediary fractions in MRC5-SV as well as CS1AN-SV cells, with different forms prevailing in each compartment (see figure 3 legend). CTSB was mainly localized in the lysosomal fraction, as expected for a lysosmal protein, but it was also detected in the mitochondrial and intermediary fractions, and this happened to a larger extent in CS1AN-SV cells. The mitochondrial localization of CTSB is in agreement with our model of CTSB leaking from lysosomes in CS cells.

We wondered whether a specific CTSB isoform is present in mitochondria. The CTSB (-2,-3) isoform contains a new N-terminal leader sequence that allows its transport to mitochondria rather than lysosomes (24). *CTSB (-2, -3)* is an alternatively spliced isoform that encodes a truncated form of the enzyme lacking the signal peptide and part of the inhibitory propeptide, and has been reported to be inactive (25). CTSB (-2, -3), which has a molecular mass of 32 kDa, may migrate in WB at the level of the CTSB upper band, *i.e.* the single-chain active CTSB form (31 kDa) (26) (*SI Appendix,* Fig. S5 A-C), and therefore go undetected. However, *CTSB (-2, -3)* transcripts are not overexpressed in most CS patient-derived fibroblasts (*SI Appendix,* Fig. S5D). Moreover, in fractionation experiments (Fig. 3C) we observed an enrichment of the lower band (double-chain active CTSB form) rather than the upper band that may co-migrate with CTSB (-2, -3). These data suggest that mitochondrial CTSB does not correspond to the (-2,- 3) isoform, which is thus unlikely responsible for POLG1 degradation, but rather to the two classic active forms of the protein. In normal conditions, these active forms are prevalently found in lysosomes, further supporting the hypothesis of lysosomal leakage of CTSB.

To assess the subcellular localization of active CTSB, we performed super-resolution structured illumination microscopy (SIM) using the Magic Red CTSB activity probe. Co-labelling of this probe with MitoTracker, a fluorescent dye that stains mitochondria in live cells, showed colocalization of CTSB activity in mitochondria in CS1AN-SV cells (Fig. 3D), in addition to extramitochondrial CTSB activity. To note, both mitochondrial and extramitochondrial CTSB activities were observed also in control (WT) MRC5-SV cells (*SI Appendix,* Fig. S5E), although the signal was low in both compartments, compatibly with lower levels of CTSB in these cells compared to CS1AN-SV cells. Thus, our results are compatible with CTSB-dependent POLG1 degradation in mitochondria.

### RAC1 is stabilized by HTRA3 and interacts with CTSB

We wondered wether HTRA3 may stablilize other proteins possibly implicated in POLG1 degradation. The RAC1 signaling pathway was found differentially enriched in a cancer related high-HTRA3 context(27), and explorative MS analyses of HTRA3- and POLG1-interactors suggested that RAC1 interacts with either proteins (not shown). Activation of the small GTPase RAC1 has been demonstrated to induce CTSB-mediated intracellular protein degradation(28). We thus hypothesized that RAC1 plays a role in the CTSB/HTRA3 containing complex that degrades POLG1 (scheme in Fig. 4A). Co-immunoprecipitation experiments showed that the active form of CTSB co-immunoprecipitates with endogenous RAC1 in CS and WT fibroblasts (Fig 4B). Moreover, HTRA3 downregulation in WT and CS cells resulted in depletion of RAC1 (Fig 4C), as it was also the case for CTSB (see Fig. 2B above), suggesting that HTRA3 also stabilizes RAC1. The regulation of RAC1 by HTRA3 is post-translational since *RAC1* transcripts increased upon HTRA3 silencing (Fig 4D), probably as a compensatory mechanism for the depleted protein, again as it was the case for CTSB. Accordingly, ectopic overexpression of HTRA3 increased the levels of RAC1 (Fig 4E, left panel). In further analogy with CTSB, treatment with the translation inhibitor cycloheximide resulted in larger amounts of RAC1 when HTRA3 was overexpressed (Fig 4E, right panel), indicating that HTRA3 stabilizes RAC1. In agreement with these findings, restoration of the HTRA3 protein content in CS1AN-SV cells 96 h post-transduction (*i.e.,* halted silencing) (see above *SI Appendix,* Fig. S4B and C) was accompanied by an increase of RAC1 protein (Fig. 4F) and transcript (Fig. 4G). Conversely, silenced MRC5-SV cells that maintained HTRA3 depletion 96 h post-transduction displayed reduced RAC1 protein (Fig. 4F) despite increased transcription (Fig. 4G), *i.e*. further indicating destabilized RAC1 upon HTRA3 depletion.

**Figure 4.**
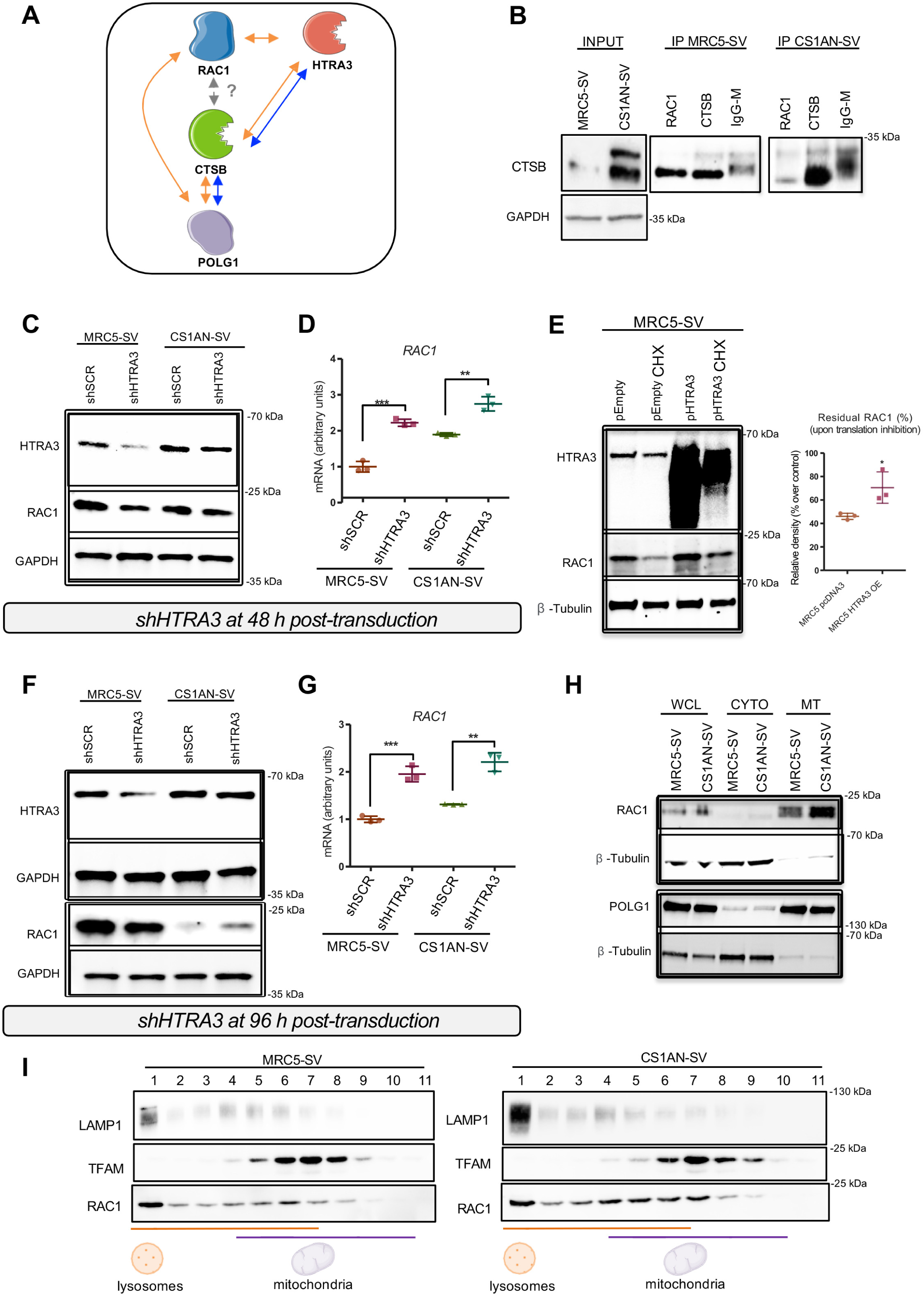
RAC1 is stabilized by HTRA3 and interacts with CTSB. (**A**) Schematic representation of protein interactions (orange arrows: interactions suggested by mass spectrometry analysis of HTRA3 and POLG1 interacting partners; blue arrows: interactions shown by co-IP; dashed grey arrows: hypothesized interaction). (**B**) Co-immunoprecipitation assays showing interaction between CTSB and HTRA3 and POLG1 proteins in WT (MRC5-SV) and CS (CS1AN-SV) cell lines. Immunoprecipitation with Ig isotypes of corresponding antibodies served as control. IP: immunoprecipitation (*same experiment as in Fig 1B*). (**C**) WB of HTRA3 and RAC1 in MRC5-SV and CS1AN-SV cells knocked-down for HTRA3 (shHTRA3) at 48 h after sh-RNA transduction (GAPDH was used as loading control) (*same experiment as* Fig. 2A and 2B). (**D**) RT-qPCR of *RAC1* at 48 h after sh-RNA transduction (**E**) WB of HTRA3 and RAC1 in MRC5-SV cells transfected with a vector coding for HTRA3 (pHTRA3) or an empty vector (pEmpty) and treated or not with 30 µg/ml of cycloheximide (CHX) for 24 h (*left panel*) (*same experiment as in* Fig. 2D). The percentages of RAC1 remaining after CHX treatment were determined by quantifying band intensities normalized to β-tubulin (loading control) and calculating the difference between treated and non-treated conditions for both the pEmpty vector and pHTRA3 (*right panel*). (**F**) WB of HTRA3 and RAC1 96 h post shHTRA3 transduction, using GAPDH as loading control (*same experiment as SI Appendix, Fig. S4B-D*). (**G**) Quantitative RT-qPCR of *RAC1* at 96 h post shHTRA3 transduction. (**H**) WB of RAC1 in whole cell lysate (WCL), subcellular mitochondrial (MITO), and cytosolic (CYTO) fractions in MRC5-SV and CS1AN-SV cells (*same experiment as SI Appendix,* Fig. 3B). β-tubulin is used as a cytosolic marker and POLG1 as a mitochondrial marker. **I**) Immunoblot analysis of RAC1 in MRC5-SV and CS1AN-SV samples fractionated on an Optiprep density gradient (*same experiment as SI Appendix,* Fig. 3C). LAMP1 is used as a lysosomal marker and TFAM as a mitochondrial marker.

Thus, CTSB and RAC1 proteins, that interact with each other as evidenced by immunoprecipitation experiments, responded similarly to changes in HTRA3 levels, suggesting that also RAC1 is stabilized by HTRA3. However, is not clear whether RAC1 is part of the complex that leads to POLG1 degradation.

### RAC1 silencing leads to a transitory downregulation of CTSB and HTRA3 proteins and degradation of a fraction of POLG1

We wondered whether RAC1, similarly to the members of the POLG1 degradation complex (CTSB and HTRA3) was present at higher levels in CS than control cells. However, no differences in the levels of RAC1 protein were observed between immortalized MRC5-SV and CS1AN-SV, as well as between primary CTL (WT) and patient-derived (CS and UVSS) fibroblasts (*SI Appendix,* Fig. S6A and B), suggesting that this protein may have a different role than HTRA3/CTSB, if any, in the homeostasis of POLG1.

Previous reports showed localization of RAC1 in mitochondrial membranes in addition to the prevalent cytosolic localization(29). We detected RAC1 to a large extent in the mitochondrial fraction, in addition to cytosolic and lysosomal fractions, in subcellular fractionation (Fig 4H) as well as OptiPrep density ultracentrifugation experiments (Fig. 4I). In both experiments the mitochondrial fraction of CS1AN-SV cells was enriched in RAC1 more than MRC5-SV cells. Thus, despite the global levels of RAC1 were similar in WT and CS cells, the mitochondrial fraction of this protein was more enriched in CS1AN-SV than MRC5-SV cells.

To assess whether RAC1 affects the proteins implicated in POLG1 depletion, we performed knock down of *RAC1* by shRNA, and this for multiple times (Fig. 5A). *RAC1* reduction of 89% in MRC5-SV and 72% in CS1AN-SV cells 36 h post-transduction (Fig. 5B, C) decreased the levels of HTRA3 and CTSB proteins in both cell types (Fig. 5C), indicating that RAC1 is necessary for the maintenance of, at least a fraction of, these two proteins. Upon *RAC1* silencing, *HTRA3* and *CTSB* transcripts increased (Fig. 5D, E), compatibly with enhanced transcription to compensate for protein depletion. Conversely, at 48 h and 96 h post-transduction, despite maintenance of RAC1 knock down (Fig. 5F, G; Fig. 5J, K, respectively), the level of HTRA3 and CTSB (Fig. 5G, K) proteins were restored in both cells, suggesting that their “stabilization” by RAC1 is only transitory, and in the long term it is either provided by other factors or is no longer necessary. Since the levels of HTRA3 and CTSB proteins were restored at these time points, their transcription returned to normal levels in MRC5-SV cells (Fig 5H, I, L, M). This was not the case in CS1AN-SV cells, where transcription of *HTRA3* and *CTSB* increased upon RAC1 silencing (Fig 5H, I, L, M), suggesting that maintenance of high levels of these proteins, as it is the case in CS1AN-SV cell, is associated to a high protein turnover. In summary, in WT and CS cells HTRA3 and CTSB behave similarly upon RAC1 depletion (*SI Appendix,* Fig. S6C and D), displaying a transitory decrease followed by restored levels that require maintained or increased transcription in MRC5-SV or CS1AN-SV cells, respectively, as the native levels of these proteins are higher in *CSB*-deficient cells.

**Figure 5.**
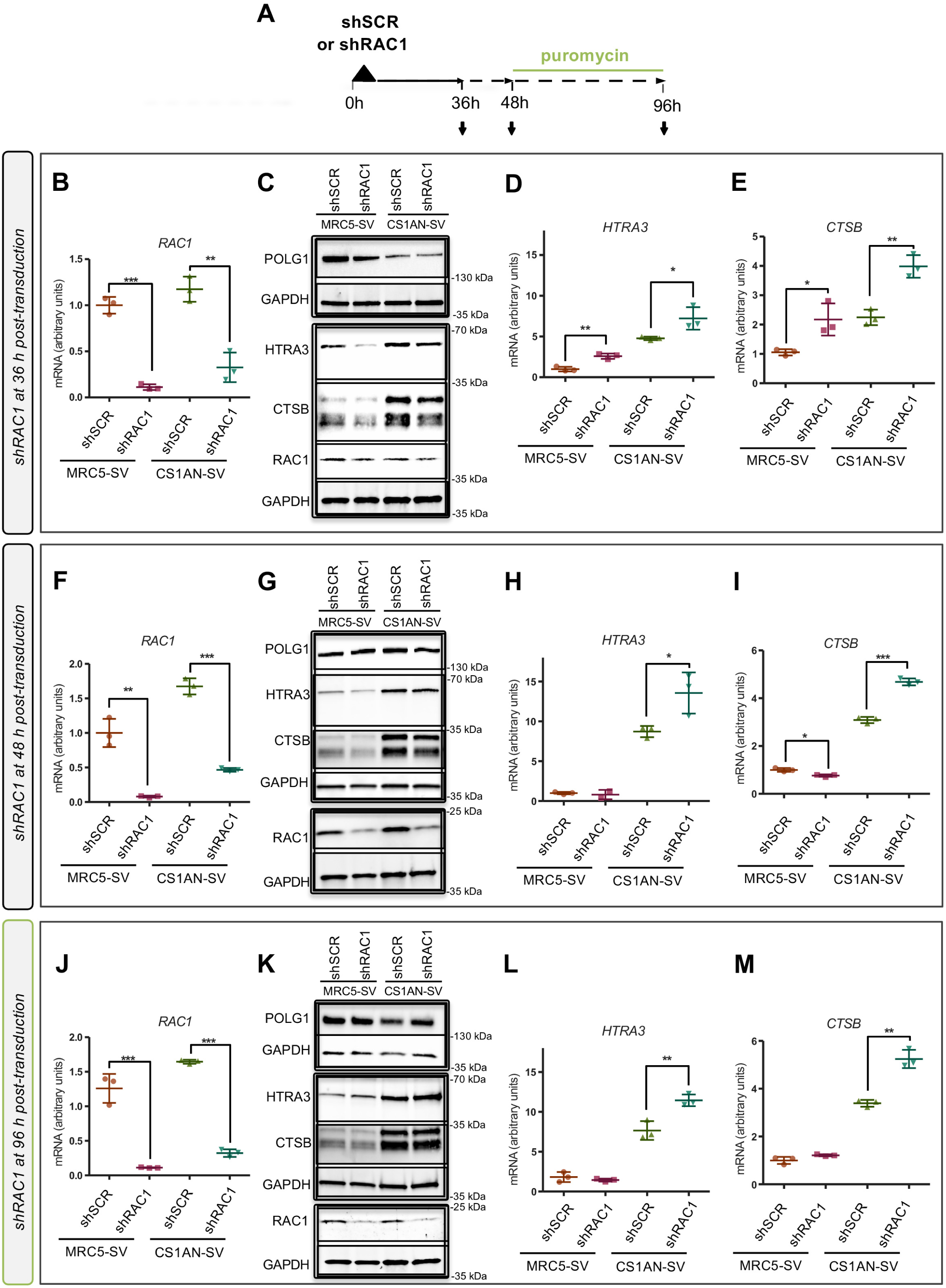
RAC1 downregulation leads to a rapidly compensated downregulation of CTSB and HTRA3 proteins. (**A**) scheme of the experiment of *RAC1* silencing (or scramble) and analysis performed at different time points: 36 h, 48 h, and 96 h post-transduction. (**B**) RT-qPCR of *RAC1* at 36 h post-transduction of sh-RNA (RAC1 or scramble) in MRC5-SV and CS1AN-SV cells. (**C**) WB analysis of POLG1, HTRA3, CTSB and RAC1 at this early timepoint, with GAPDH as a loading control. RT-qPCR of (**D**) *HTRA3* and (**E**) *CTSB* at the same timepoint as previous panels. (**F**) Quantitative RT-qPCR of *RAC1* in MRC5-SV and CS1AN-SV fibroblasts at 48 h after transduction with shRAC1 or shSCR. (**G**) WB of POLG1, HTRA3, CTSB, and RAC1 at 48 h after transduction (each with the loading control GAPDH). RT-qPCR of (**H**) *HTRA3* and (**I**) *CTSB* at 48 h post-transduction. (**J**) RT-qPCR of *RAC1* at 96 h following transduction of sh-RNA (**K**) WB of POLG1, HTRA3, CTSB, and RAC1 at this timepoint, using GAPDH as loading control. RT-qPCR of (**L**) *HTRA3* and (**M**) *CTSB* also at day 4 post-transduction. To note, RAC1 silencing had the largest effect 36h post-shRNA transduction, reducing the HTRA3 and CTSB protein content, which was then rapidly restored (48h post-transduction) in both cells. Samples on the same blot are framed; each frame displays the respective GAPDH used as loading control. RT-qPCR: n=3 independent experiments; mean ± SD. \**p* ≤ 0.05, \*\**p* ≤ 0.01, \*\*\**p* ≤ 0.001; based on unpaired t test comparisons to respective shSCR.

We wondered whether the effect of *RAC1* silencing on proteins involved in POLG1 degradation translated into changes in POLG1 levels. In CS1AN-SV cells 36 h post-transduction (silencing), despite reduced levels of HTRA3 and CTSB, the POLG1 content did not change (*SI Appendix,* Fig. 5C and E), suggesting that either i) residual HTRA3/CTSB in these cells are sufficient to maintain POLG1 degradation, or ii) RAC1 is not necessary for POLG1 degradation in these cells, or else iii) RAC1 enhances POLG1 stability and counteracts HTRA3-CTSB degradation. The first two hypotheses are not compatible with MRC5-SV cells that carry a natively high POLG1 content. Indeed, contrary to either of these two hypotheses, in MRC5-SV cells POLG1 levels decreased upon *RAC1* silencing (36h). Conversely, the hypothesis (iii) is compatible with RAC1 interacting with or sequestering CTSB (and/or HTRA3), or at least a fraction of, and by this hindering its/their capacity to degrade POLG1. The larger mitochondrial localization of RAC1 in CS1AN-SV than MRC5-SV cells, and the prevalent mitochondrial localization of POLG1 in both cells, may explain a different access of RAC1 to HTRA3/CTSB in mitochondria, and thereby POLG1 degradation in these two cell types. To note, at 48 h and 96 h post-transduction (*RAC1* silencing), the levels of POLG1 protein no longer decreased in MRC5-SV cells (Fig. 5G,K, Fig S6F,G), suggesting that RAC1-free HTRA3/CTSB (proteolytic) complexes are no longer available to interact with POLG1 (scheme in *SI Appendix,* Fig. S6H). Thus, RAC1 appears to participate in the homeostasis of POLG1 by modulating its HTRA3/CTSB proteolytic effectors, probably depending on its mitochondrial *versus* non-mitochondrial localization.

### CTSB overexpression is recapitulated in cellular senescence

We reported that HTRA3 upregulation and subsequent POLG1 depletion identified in fibroblasts of progeroid CS patients are recapitulated during replicative senescence of normal cells, and this process is triggered by progressive depletion of CSB, the protein mutated in CS(30). We thus examined whether CTSB expression was upregulated in senescent fibroblasts. For that, DNA damage-dependent senescence was triggered by irradiating (10 Gy) early-passage normal primary BJ fibroblasts. High levels of the senescence marker p21 indicated that the cell population was senescent at 10 days post-irradiation and, as expected, CSB decreased and HTRA3 levels increased in senescent BJ cells (Fig. 6A), in agreement with our previous findings(30). Importantly, as predicted by the analogy between CS and senescent normal cells, high HTRA3 levels were accompanied by an increase in the CTSB protein content in BJ (Fig. 6A) as well as in IMR-90 irradiated primary fibroblasts (*SI Appendix,* Fig. S7A, senescence assessed by increased p21 levels in Fig. S7B).

**Figure 6.**
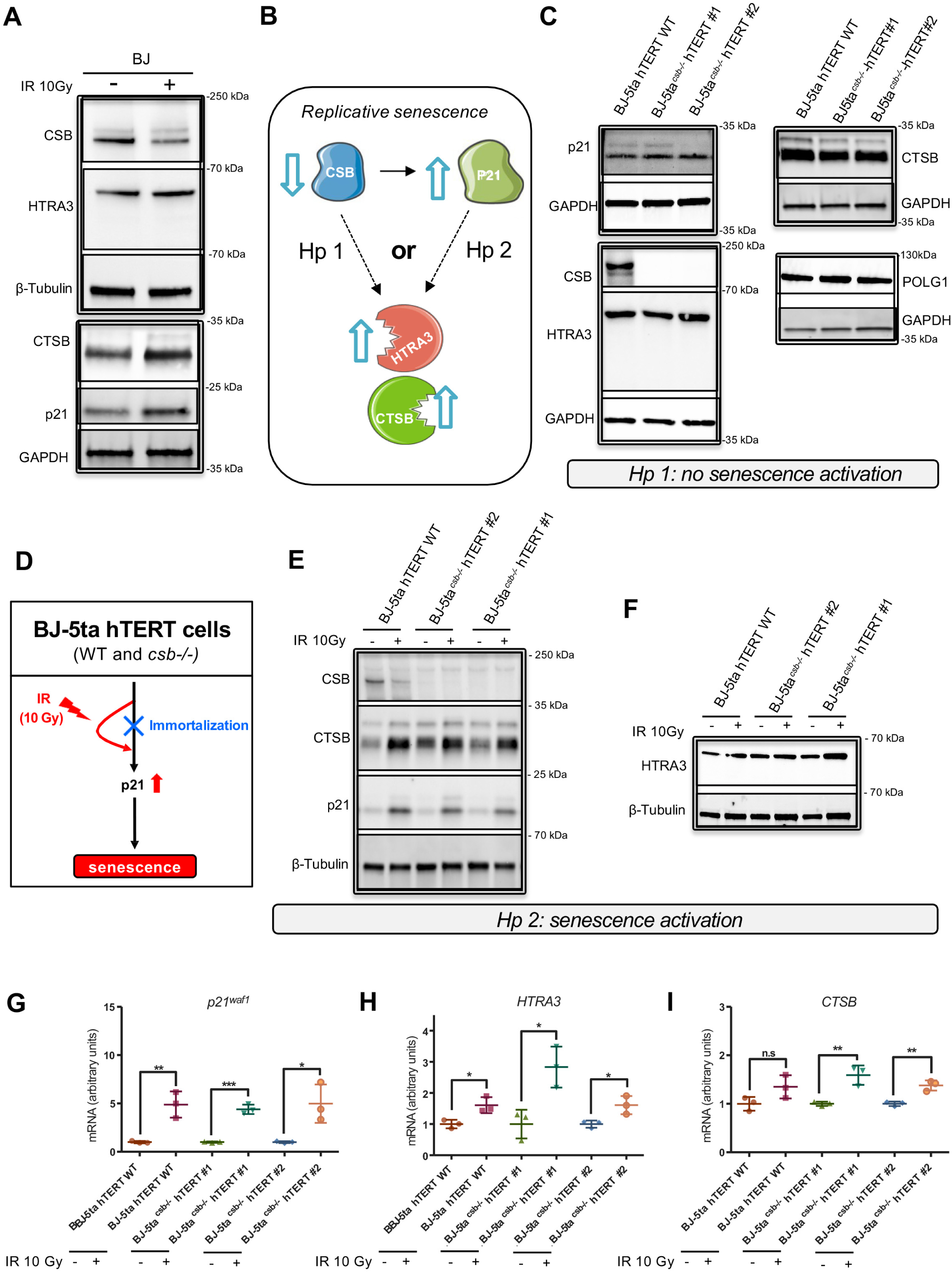
CTSB overexpression is recapitulated in cellular senescence. **(A)** WB of CSB, HTRA3 (left panel), and CTSB, and p21 (right panel) in BJ fibroblasts untreated and 10 days post-irradiation using ß-tubulin and GAPDH, respectively, as loading control. **(B)** Scheme showing the two hypotheses for the mechanism leading to senescence-related HTRA3 and CTSB overexpression. **(C)** Immunoblots of p21, CSB, HTRA3, CTSB, and POLG1 in BJ-5ta hTERT WT fibroblasts and 2 CRISPR-edited CSB knocked-out clones (BJ-5ta^csb-/-^hTERT #1 and BJ-5ta^csb-/-^hTERT #2). Frames show samples on the same blot; each displaying the respective GAPDH used as loading control. **(D)** Scheme indicating normally blocked senescence in BJ-5ta hTERT cells and activation of senescence upon irradiation at 10 Gy. WB analysis of **(E)** CSB, CTSB, p21, and **(F)** HTRA3 three days post-irradiation in BJ-5ta hTERT WT fibroblasts and BJ-5ta hTERT CSB knocked-out fibroblasts (β-tubulin was used as loading control). RT-qPCR of (**G**) *p21^Waf1^* and (**H**) *HTRA3* and (**I**) *CTSB* in the same experiment and cells schematized in panel D. Samples on the same blot are framed; each frame displays the respective GAPDH or β-tubulin used as loading control. RT-qPCR: n=3 independent experiments; mean ± SD. \**p* ≤ 0.05, \*\**p* ≤ 0.01, \*\*\**p* ≤ 0.001; based on unpaired t test comparisons to respective non-irradiated control.

The mechanism leading to senescence-related HTRA3 accumulation, and now also to CTSB accumulation, remains to be elucidated. Transcriptional regulation of HTRA3 and CTSB either directly depends on CSB that is also a transcription factor (hypothesis 1; Fig. 6B) or it is triggered by senescence effectors, and therefore by the senescence process itself (hypothesis 2; Fig. 6B). Among senescence effectors we considered p21 that we previously reported to be transcriptionally activated by decreased CSB levels (3). To disentangle these two hypotheses, it was necessary to uncouple CSB depletion from cellular senescence. For that, we knocked out CSB using CRISPR-Cas9 editing in an *ad hoc* cellular model where senescence is normally blocked, *i.e.* in immortalized cells, but can be reactivated upon specific stimuli (31). To remain as close as possible to the primary cell phenotype, we edited hTERT-immortalized fibroblasts (BJ-5ta hTERT) by inserting two alternative point mutations in *CSB*. These two independent clones have been constructed to mimic *CSB* mutations reported in CS patients. The mutations and sequencing of the two clones (BJ-5ta*^csb-/-^*hTERT #1 and BJ-5ta*^csb-/-^*hTERT #2) are detailed in Figure S8, and CSB depletion was confirmed by WB (Fig. 6C). As anticipated, knock-out of CSB that should trigger p21 expression in normal cells(3), did not affect p21 expression in this immortalized cell line (Fig 6C). Importantly, CSB depletion did not lead either to HTRA3 or CTSB accumulation (Fig. 6C), indicating that the levels of HTRA3 and CTSB proteins are not directly regulated by CSB. These experiments suggest that HTRA3 and CTSB are rather regulated by the senescence process itself. If this were the case, reactivation of senescence in CSB-depleted BJ5ta-hTERT cells (upon stress induction like irradiation (31), schematized in Fig. 6D), should lead to upregulation of HTRA3 and CTSB expression.

Senescence was observed in BJ5ta^csb-/-^ hTERT cells already 3 days after irradiation by upregulation of the senescence marker p21 (Fig. 6E, G) and the appearance of senescence-associated beta-galactosidase staining (SA-β-gal) that further increased 7 days post-irradiation (*SI Appendix,* Fig. S9). Senescence resulted in high levels of the CTSB (Fig. 6E) and HTRA3 (Fig. 6F) proteins and their transcripts (Fig. 6H, I), as predicted by our model.

These data are compatible with accumulation of the HTRA3 and CTSB proteases being dependent on the senescence process rather than CSB depletion, as anticipated in hypothesis 2. In WT (CSB-proficient) BJ5ta-hTERT cells, induction of senescence, which was confirmed by high p21 levels (Fig. 6E, G), was associated with reduced levels of CSB, in agreement with our previously findings(3). Importantly, control BJ5ta-hTERT cells also displayed increased levels of the HTRA3 and CTSB proteins and transcripts (Fig. 6E, F, H, I), further supporting the role of senescence rather than CSB levels in promoting high levels of these two proteases.

## Discussion

Mitochondrial protein homeostasis is maintained through a variety of mechanisms that overall control their synthesis, transport to the specific compartment within the organelle, and eventual degradation, processes that are closely tied to the localization of mitochondrial proteins(32). Misfolded or unassembled proteins, as well as native proteins are selectively degraded to adjust mitochondrial function to the energetic needs of the cell. This regulation becomes impaired with ageing, contributing to mitochondrial dysfunction, an hallmark of the ageing process. Mitochondrial proteases play a critical role in this quality control, ensuring proper proteins turnover (32). Despite its pivotal importance in mtDNA replication and overall, mitochondrial and cellular function, the mechanism governing POLG1 degradation have remained largely unexplored. Our study identifies a specific pathway responsible for POLG1 degradation, leading to its depletion, under progeroid and senescent conditions. Our findings show a selective POLG1 regulation by the CTSB/HTRA3 complex, which is itself regulated, at least to some extent, by the small GTPase RAC1. We previously reported POLG1 depletion in cells from patients with progeroid Cockayne syndrome (CS), that required HTRA3 overexpression (11), and was recapitulated in replicative senescence (3), a major hallmark of ageing, indicating that POLG1 depletion has links with pathophysiological ageing.

We show here that HTRA3-dependent POLG1 degradation is more complex than a direct protease/target interaction, and rather requires the implication of the HTRA3 protease-chaperone that stabilizes another protease, CTSB, that is in turn responsible for cleaving the target POLG1. This is a different type of interaction compared to known mitochondrial matrix proteases, which also act as chaperone-proteases: for instance, the Lon protease carries the protease and the chaperone activity in the same polypeptide, whereas the ClpXP protease is composed by the ClpX chaperone and the ClpP peptidase (33). Conversely, the HTRA3/CTSB partnership is specific in that despite its dual function, HTRA3 acts essentially as a chaperone and not as a protease. Moreover, neither HTRA3 nor CTSB are prevalently or exclusively located in mitochondria, as it is the case for the abovementioned matrix proteases. Indeed, HTRA3 is present in the mitochondrial as well as non-mitochondrial compartment of the cytoplasm, as we previously reported (3). Here, we demonstrate the localization of active forms of CTSB and, importantly, CTSB activity in mitochondria, with a larger mitochondrial localization in CS cells. The CTSB(-2,-3) isoform that was previously reported in mitochondria (24) is catalytically inactive and cannot properly fold into an active protease (25). Lysosomes, that contain most cellular and active forms of CSTB in normal conditions, have been reported to be dysfunctional in CS cells, due to ruptured membranes(20). We therefore hypothesized that active CTSB leaking from lysosomes is targeted to mitochondria where it degrades POLG1. In line with this hypothesis, induced disruption of the lysosomal membranes with LLOMe resulted in decrease of POLG1, both in WT and CS cells. This degradation activity seems specific to POLG1, since upon lysosome leakage we did not observe depletion of TFAM, another mitochondrial matrix protein as POLG1. To note, it was proposed that TFAM, that is normally degraded by the LON protease (34), is degraded by CTSB in aged microglia (35), suggesting that under other conditions also TFAM homeostasis is possibly regulated by CTSB. The HTRA3/CTSB interaction is favored in CS cells, where both HTRA3 and CTSB are overexpressed, and these proteins are also enriched in the mitochondrial fraction where the target POLG1 is mainly located, resulting in a larger degradation of POLG1 than in WT cells.

Although it is generally accepted that cysteine cathepsins are active only at slightly acidic pH as in the endolysosomal vesicles, actually cathepsins retain extralysosomal activity for a limited amount of time. Similarly, CTSB has been reported to be highly active in a pH range of 7.5–8 (36), which is compatible with the pH of the mitochondrial matrix (pH 7.7–8.2) (37-39). However, the enzyme is more unstable and spontaneously denatures at this high pH (40), reinforcing the notion of the requirement of a chaperone like HTRA3 to remain active. In agreement with this notion, HTRA3 was shown to protect *in vitro* the citrate synthase against thermal inactivation in a concentration-dependent manner (22).

We also identified the small GTPase RAC1 as a regulator of HTRA3 and CTSB and, ultimately, POLG1 degradation. RAC1 responds as CTSB upon HTRA3 downregulation and overexpression, suggesting that also RAC1 is stabilized by HTRA3. Subcellular fractionation analysis demonstrated the mitochondrial localization of the CTSB partners RAC1 and HTRA3, in line with previous reports by our and other labs (3, 41, 42), and at a higher extent in CS than WT cells. Silencing of RAC1 transitorily stabilizes HTRA3 and CTSB but by this way blocks them from digesting POLG1. It is possible that RAC1 alters the properties of HTRA3 and CTSB (or their complex), as it is the case for components of the NADPH oxidase complex, where RAC1 induces conformational changes in the p67phox protein that allow its binding to gp91phox; an interacting complex that is required for oxidative burst (43). Altogether, the demonstrated interaction between RAC1 and CTSB, the correlative levels of HTRA3 and CTSB, their mitochondrial localization, and the mitochondrial localization of RAC1 are compatible with RAC1-free HTRA3/CTSB digestion of POLG1 (scheme in *SI Appendix,* Fig. S6H). Therefore, RAC1 counteracts POLG1 degradation performed by HTRA3/CTSB. All these processes occur at a larger extent in CS than WT cells, resulting in more extensive POLG1 degradation in the former than in the latter. To note, mitochondrial RAC1 operates on available levels of HTRA3/CTSB present in the organelle, that are larger in CS cells. It is unclear whether RAC1 also interacts with POLG1, as suggested by MS experiments, because POLG1 and RAC1 Co-IP experiments were not feasible with the available antibodies. Since RAC1 is implicated in multiple functions in mitochondria, the mitochondrial relocalization of RAC1 in CS cells may have implications beyond the participation in POLG1 degradation. For instance, the interaction of RAC1 with BCL2 in mitochondria stabilizes the BCL2-mediated superoxide production that is required to maintain a mild pro-oxidant intracellular level, resulting either in a cyto-protective or cytotoxic effect, depending on cell type and environmental conditions (42).

POLG1 degradation in CS cells, that we showed affecting mitochondrial function, seems thus linked to the progeroid and neurodegenerative phenotype of this syndrome (POLG1 degradation was not reported in UVSS cells, derived from patients that age normally and do not display neurodegeneration) (11). Likewise, several mitochondrial defects have been implicated in the pathogenesis of CS, including mitochondrial membrane depolarization, increased mitochondrial content due to defective mitophagy, increased ROS production and increased membrane potential (44-46). The mitochondrial involvement in CS pathogenesis is also supported by the stricking similarity of the CS phenotype to typical aspects of mitochondrial diseases, such as delayed-onset and progressive course, tissue-specific manifestations, and variation in individual predisposition to degenerative defects (19, 47). This is particularly interesting for a progeroid disease since decline in mitochondrial function is a major hallmark of ageing (15). In this context, we showed that POLG1 degradation is not limited to cells of progeroid CS patients, but it is also recapitulated during replicative senescence of normal cells. In these cells, progressive depletion of CSB (the protein that is impaired in CS) triggers senescence by exposing the p21 promoter that can be thus activated to prime the program of replicative senescence(3).

We show here the relevance of the CTSB/HTRA3-dependent POLG1 degradation in normal cells and disentangle its regulation upon replicative senescence rather than CSB depletion. First, we show that CTSB specifically accumulates in CS cells compared to WT or UVSS (*CSA* mutated but not progeroid), with a possible implication of CTSB in the CS progeroid phenotype. In this context, CTSB overexpression had been linked to regular ageing and neurodegeneration in the mouse hippocampus. Moreover, CTSB-deficient mice display reduced increase of age-dependent oxidation and inflammation (35). Similarly, a direct correlation between serum CTSB concentration and the age-related decline in cardiovascular and renal functions has been reported(48). In our model, and again similarly to HTRA3 (3), we observed a CTSB accumulation during DDR-dependent senescence, a process linked to ageing, suggesting a tight regulatory link between these two proteins. This last observation is in agreement with a previous study showing that CTSB is a member of the senescence-associated secretory phenotype (SASP) (49).

Second, we show that induction of HTRA3 and CTSB is not directly dependent on CSB levels (as it is the case for instance for the cell cycle regulator p21(3)) but rather on cellular senescence itself. Indeed, we have established a cellular system that is able to uncouple CSB depletion from cellular senescence, two processes that are normally linked (decreased CSB levels in primary cells leads to p21 upregulation and senescence)(3). For that, we knocked-out CSB with CRISPR-Cas9 in immortalized human BJ5ta fibroblasts that do not undergo senescence, unless properly activated. We discovered that CSB ablation does not lead to changes in HTRA3 or CTSB levels nor, consequently, POLG1 depletion, revealing that the expression of the two proteases is not directly regulated by CSB. Exogenous stress-induction of senescence, which can be exceptionally re-activated in these immortalized cells, was sufficient to trigger HTRA3 and CTSB overexpression, demonstrating that their regulation is linked to the senescence process. This result suggests that CS-specific POLG1 depletion and the consequent mitochondrial defects, that are specifically linked to the progeroid phenotype (2), are not directly dependent on CSB impairment itself but on the induction of senescence that results from it. Thus, we show a DNA repair-independent mechanistic model for POLG1 degradation that contributes to POLG1 homeostasis in basal conditions, and is hyperactivated in CS and cellular senescence.

In conclusion, the present study uncovers a mechanism of mitochondrial POLG1 degradation driven by the interaction between the serine protease HTRA3, which acts as a chaperone to stabilize the cysteine protease Cathepsin B, that reaches mitochondria following destabilization of lysosomal membranes. This HTRA3/CTSB-dependent degradation of POLG1 is partially counteracted by the GTPase RAC1, further regulating POLG1 degradation. This mechanism is particularly relevant in cells from progeroid Cockayne syndrome as well as senescent cells, where POLG1 depletion occurs, linking it directly to processes of pathophysiological ageing. Notably, it is the process of senescence itself, rather than the initial CSB impairment, that drives POLG1 degradation, further highlighting its significance in ageing biology.

## Materials and methods

### Cells and cell culture

Patient fibroblasts were obtained from skin biopsies excised by dermatologists from unexposed body sites. Patients (or parents or legal guardians) provided informed consent to receive diagnosis and genetic analysis. This study was approved by the French Ministry for Research (project MolDefCS 2003-007, DC-2023-5868).

Human primary skin fibroblasts from healthy individuals, UVSS and CS patients, MRC5-SV and CS1AN-SV SV40-transformed cell lines (a generous gift from A. R. Lehmann, Sussex University, Sussex, United Kingdom), were cultured with Dulbecco’s modified eagle medium (DMEM; Gibco), supplemented with 2mM L-glutamine (GlutaMAX; Gibco), 10% foetal bovine serum (FBS; Gibco) and 1% Penicillin-streptomycin (Gibco). Normal female human foetal lung IMR-90 fibroblasts (ATCC; CCL-186) and BJ skin fibroblasts (ATCC; CRL-2522) were cultured in minimum essential medium (MEM, Gibco) supplemented with 2 mM L-glutamin (GlutMAX), 10% FBS, 1% Penicillin–streptomycin, 1% nonessential amino acids (Gibco) and 1% sodium pyruvate (Gibco) BJ-5ta hTERT immortalized human fibroblasts (ATCC; CCL-186) cultured in a 4:1 mixture of DMEM and Medium 199 (Gibco) supplemented with 10% FBS and1% Penicillin–streptomycin. All cells were cultured in 20% O_2_/5% CO_2_ at 37 °C.

When indicated, MRC5-SV and CS1AN-SV were treated for 3 h with 250 µM and 1000 µM of L-leucyl-L-leucine methyl ester (LLOMe) (Cayman Chemical, 6491834; diluted in DMSO) followed by 1h of drug withdrawal, or with 30 µg/ml cycloheximide (CHX) (Clinisciences, A8244; diluted in DMSO) for 24h. Control experiments were conducted in the presence of an equivalent amount of DMSO.

For radiation-induced senescence, confluent cells were irradiated at 10 Gy with the Xstrahl RS320 irradiator (X-ray). Cell culture medium was refreshed after irradiation and cells were incubated overnight. The following day, cells were diluted 1:3 and cultured for the indicated times.

### shRNA cloning, transfection and transduction

Lentivirus particles were generated from the shRNA plasmids or scrambled (SCR) shRNA for control (see *SI Appendix,*Table S1). shRNA plasmids, carrying ampicilin resistance, were amplified in *E. coli* grown overnight in Luria Bertani (LB) broth medium with 100 µg/ml ampicillin, and extracted and concentrated using the NulceoBond® Xtra Maxi Plus kit, Plasmid DNA purification kit (Macherey-Nagel, 740416.10) following the manufacturer’s instructions. Purified shRNA plasmids were co-transfected with psPAX2 (packaging) (Addgene, 12260) and pMD2.G (envelope) (Addgene, 12259) plasmid in HEK-293T cells, using calcium phosphate transfection method (50). Forty-eight hours post-transfection the culture medium was collected, filtered through 0.45 µm filters and concentrated by ultra-centrifugation at 19,000 rpm for 90 min at 4 °C. The pellet resulting from this centrifugation was resuspended to a final volume of 250 µl in phosphate buffered saline (PBS).

Transduction of MRC5-SV and CS1AN-SV fibroblasts with shSCR, and shRNA (shCTSB/shHTRA3/shRAC1) was performed by incubating cells with the viral particles in the presence of 8 µg/ml of Polybrene, which acts as a transduction enhancer (Sigma-Aldrich, H9268). After the overnight incubation, medium containing the viral particles and Polybrene was removed. When indicated cells were collected at 48 h post-transduction. Alternatively, cells were selected with 1 µg/ml puromycin at 48 h post-transduction and collected at the indicated points.

### Western blotting

Cells were lysed for 30 min on ice with lysis buffer (1 mM EDTA, 50 mM Tris-HCl pH 7.5, 150 mM NaCl, 1% Triton X-100, 0.1% SDS and protease/phosphatase inhibitor mixture (Roche)). The whole cell extracts were sonicated (Bioruptor, Diagenode) 5 times for 10 sec, with 10 sec intervals, at medium frequency at 4 °C. Protein concentration was determined with the Bradford assay, and samples were boiled 5 min at 95 °C in the presence of NuPAGE LDS sample buffer (Invitrogen NP0007) and NuPAGE Sample Reducing Agent (Invitrogen NU0004). 10 µg of protein were loaded for SDS/ PAGE (4-12% Bis-Tris Gel, Thermo-Fisher Scientific or 4-15% Mini-PROTEAN TGX gel, Bio-Rad) and transferred on a nitrocellulose transfer membrane by Trans-Blot Turbo system (Bio-Rad). The membrane was then blocked with 5% nonfat dry milk in 1x PBS with 0.1% Tween20 (Sigma-Aldrich, P1379) for 1h at room temperature (RT). Primary antibodies at indicated concentrations were incubated in 1% nonfat dry milk in 1x PBS overnight at 4°C. The following primary antibodies were utilized at the specified dilutions: rabbit polyclonal α-HTRA3 (1:1000; Sigma-Aldrich, HPA021187), mouse monoclonal α-CTSB (1:1000; Abcam, ab58802), mouse monoclonal α-RAC1 (1:1000; Abcam, ab33186), rabbit polyclonal α-CSB (1:500; Abcam, ab96089), rabbit polyclonal α-POLG1 (1:1000; Abcam, ab128862), rabbit polyclonal α-TFAM (1:1000; Sigma-Aldrich, HPA040648), rabbit polyclonal α-LAMP1 (1:1000; Abcam, ab24170), and mouse monoclonal α-p21 (1:200; BD Biosciences, AB396414). After five washes of 5 min in PBS containing 0.1% Tween-20, the membrane was incubated with the secondary antibodies (hFAB Rhodamine α-GAPDH (1:5000; Bio-Rad, 12004168), StarBright Blue 700 Goat α-Mouse IgG (1:5000; Bio-Rad, 12004158), StarBright Blue 700 Goat α-Rabbit IgG (1:5000; Bio-Rad, 12004162), CF770 Goat α-Mouse IgG (1:20000; Biotium, 20070), CF770 Goat α-Rabbit IgG (1:20000; Biotium, 20078), HRP Goat α-Mouse IgG (1:10000; Thermo Fisher Scientific, 31430), HRP Goat α-Rabbit IgG (1:10000; Thermo Fisher Scientific, 31460) for 1h at RT. After five washes of 5 min in PBS containing 0.1% Tween-20, the membrane was revealed by chemiluminescence or fluorescence using a Chemidoc™ MP imaging system (Bio-Rad). ATX Ponceau S Red (Sigma-Aldrich, 09189) was employed as a further marker of protein content. Quantification of Western blot bands was performed using the ImageJ software.

### Immunostaining

Cells were plated on glass slides, fixed in 4% paraformaldehyde (PFA) for 15 min at RT and permeabilized with 0.5 % Triton X-100 (Sigma-Aldrich, T9284) for 15 min at RT. Permeabilized samples were incubated in blocking buffer (5% BSA (Gerbu Biotechnik, 1503) in PBS) for 1h at RT. Primary antibodies were incubated overnight at 4°C in a humidified chamber as follows: mouse monoclonal α-CTSB (1:200; Abcam, ab58802), rabbit polyclonal anti α-POLG1 (1:200; Abcam, ab128862), and rabbit polyclonal α-LAMP1 (1:1000; Abcam, ab24170). Secondary antibodies used were Alexa Fluor 488 α-mouse IgG (1:2000; Thermo Fisher Scientific, A-11001), Alexa Fluor 488 α-rabbit IgG (1:2000; Thermo Fisher Scientific, A-11008), Alexa Fluor 555 α-mouse IgG (1:2000; Thermo Fisher Scientific, A-31570), Alexa Fluor 555 α-rabbit IgG (1:2000; Thermo Fisher Scientific, A-21428), and 5 µg/ml Hoechst 33342 (Sigma, 14533) for nuclear counterstaining, all incubated for 1h at RT.

### CTSB activity

CTSB enzymatic activity was detected by Magic Red reagent (Abcam, ab270772) according to manufacturer’s protocol. Briefly, 20 µl of a reconstituted Magic Red staining solution was added to 280 µl medium and cells were incubated in the mixture at 37 °C for 30–60 min protected from light. Cells were then subjected to immunostaining and imaging procedures previously indicated.

### Confocal acquisition and quantification

Cells were imaged using a Nikon Ti2E confocal fluorescent microscope with Yokogawa CSU-W1 Spinning Disk integrated with an 63x/1.4 oil objective and a Hamamatsu Orca Flash 4 camera (UTechS PBI of Institut Pasteur, Paris). For 3D analysis, 25–38 Z-stack images covering the whole depth of the cell were acquired. Quantifications were done with the ImageJ 2.0 v software (NIH) by dividing the sum of the gray values of all the pixels in each cell by the number of pixels per cell and cell surface. Images are shown as summed projections of several confocal planes. For each condition, 30–60 cells were analyzed from three independent experiments.

### Structured illumination microscopy (SIM)

MRC5-SV and CS1AN-SV were plated on precision glass coverslips thickness No. 1.5 H (Marienfeld Superior). Next day, cells were stained with stained with MitoTracker Deep Red and CTSB Magic red. For MitoTracker staining cells were incubated 1 h with 200 nM MitoTracker Red FM (Invitrogen). For CTSB Magic red staining see specific section. Immunofluorescence was performed as indicated in the previous section, using Fluoromont-G (Thermo fisher Scientific) as mounting medium. The images were obtained using a Zeiss LSM 780 Elyra PS1 microscope (Carl Zeiss, Germany) using a 63X/1.4 oil Plan Apo objective with a 1.518 refractive index oil (Carl Zeiss) and an EMCCD Andor Ixon 887 1 K camera for the detection. Fifteen images per plane per channel (five phases, three angles) were acquired with a Z-distance of 0.2 µm to perform 3D-SIM images. To optimize the signal to noise ratio, acquisition parameters were adapted from one image to one other. The ZEN software was used to process and align the SIM images. The final rendering of the images between control and treated cells corresponds to different brightness/contrast treatments. Image acquisition and processing was done by Audrey Salles (UTechS Photonic BioImaging PBI (Imagopole), Institut Pasteur).

### Subcellular Fractionation Assays

Subcellular fractionation assays were mostly performed as previously reported(51). Briefly, four 150mm dishes, at 80% confluence, were washed twice with PBS, detached using a cell scraper and pelleted by centrifugation at ∼450 × g for 5 min. The cell pellet was resuspended in mitochondria isolation buffer (MIB, 10 mM Tris/MOPS, 200 mM sucrose, 1 mM EGTA/Tris and protease inhibitors), and then lysed using y 10 strokes of a 28-gauge syringe. Lysis efficiency was confirmed by visualizing 10 µl of cell lysate stained with 0.5% trypan blue under a light microscope. As whole cell lysate, an aliquot of the obtained cell lysis was utilized. The rest of the cell lysate was centrifuged at 600 g for 10 min at 4 °C to separate the nuclei, that remained in the pellet. To remove any nuclei contaminants that could remain, the supernatant was collected and centrifuged again at 600 g for 10 min at 4 °C. To pellet mitochondria, the supernatant was centrifuged at 7000 g for 10 min at 4 °C. The pellet, washed once with MIB and resuspended in NuPAGE LDS buffer represented the mitochondrial fraction. The supernatant constituted the cytosolic fraction, and was centrifuged at 7000 × g for 10 min at 4 °C to remove remaining mitochondrial contamination, and finally diluted in NuPAGE LDS 2X (CYTO).

### Co-Immunoprecipitation

Co-Immunoprecipitation (co-IP) was performed using a Dynabeads™ Protein G (Thermo Scientific, 10003D). The Dynabeads-Ab complex was prepared by incubating with 5ug of each specific antibody (against RAC1 (Abcam, 33186), CTSB (Abcam, 58802), HTRA3 (Sigma-Aldrich, HPA021187, POLG1 (Abcam, 128862) with Dynabeads™ Protein G (previously washed once with PBS and lysis buffer and resuspended in lysis buffer) with rotation overnight at 4 °C. Irrelevant normal mouse or rabbit IgGs were used as control (SC-2025 or SC-2027 respectively, Santa Cruz). The following day, cells washed twice with PBS and collected using a cell scraper. Cells were lysed for 30 min on ice on lysis buffer (1 mM EDTA, 50 mM Tris-HCl pH 7.5, 150 mM NaCl, 1% Triton X-100, 0.1% SDS and protease inhibitor mixture (Roche)). 1 mg of cell lysate was added to the Dynabeads-Ab complex and the mix as incubated with rotation overnight at 4 °C, to obtain the Dynabeads-Ab-antigen complex. The immunocomplexes were then washed twice with lysis buffer and eluted from from Dynabeads Protein G by boiling 10 min at 95 °C in 2X NuPAGE LDS sample buffer and NuPAGE Sample Reducing Agent and removing the Dynabeads Protein G with a magnet, being subsequently analysed by Western blotting using the procedure previously specified. To eliminate interference from the denatured/reduced heavy and light chains of the IP antibody, HRP–conjugated secondary antibodies able to detect only native IgG were used (1:200; Rockland, TidyBlot, 18-8817-30).

### OptiPrep Gradient Fractionation

OptiPrep gradient fractionation was performed as described previously(23). Briefly, cells were washed twice with PBS, harvested with a cell scraper and pelleted by centrifugation at ∼450 × g for 5 min. A lysosome enrichment kit (Pierce, Cat #89839, USA) was used to perform subcellular fractionation. Pellets were resuspended in Lysosome Enrichment Reagent (LER) “A” buffer containing 1% protease inhibitors and incubated for 2 min on ice. The cell suspension was homogenized on ice using a ball-bearing cell homogenizer (Isobiotec). Lysis efficiency was confirmed by visualizing under a light microscope 10 ul of cell lysate stained with 0.5% trypan blue. After 15 passages through the homogenizer, 95% of cell membrane breakage was achieved. LER “B” containing 1% protease inhibitors was then added to the lysed cells. Cell nuclei and membranous debris were pelleted by centrifugation at 600 × g for 10 min at 4 °C and the postnuclear supernatant was retained.

To achieve a discontinuous density gradient, gradient solutions (30%, 27%, 23%, 20% and 17%) were prepared by mixing the OptiPrep medium (supplied as a 60% solution in H2O) with gradient dilution buffer (mixture of LER A and LER B 1:1), and gradient solutions were carefully overlaid in a 5 ml ultracentrifuge tube (Beckman, 326819) in descending concentration order. The postnuclear supernatant sample was mixed with OptiPrep medium to a final concentration of 15% OptiPrep, and was placed on top of the density gradient and centrifuged at 145.000 × g for 4h at 4 °C using an Optima MAX-XP ultracentrifuge (Beckman Coulter) and a swinging bucket Rotor MLS-50. After centrifugation, 11 fractions were carefully collected from the top of the gradient. To separate the lysosomes and mitochondria from the cytosol, isolated fractions were mixed with 1ml of PBS and centrifuged at 20,000 × g for 30 min at 4 °C.

The pellets were resuspended in 400 ul of PBS and constituted the lysosomal and mitochondrial fractions. These fractions were used for Western blotting analysis for the identification of organelle markers and proteins of interest.

### SA- β-gal staining

Cultured cells were washed in PBS and fixed at room temperature (RT) for 5 min with 2% paraformaldehyde (PFA, Alfa Aesar, 43368) and 0.2% glutaraldehyde (Sigma-Aldrich, G6257) in PBS (Gibco) and rinsed twice with PBS. Fixed cells were stained at 37 °C overnight with 1 mg/ml X-Gal solution (Thermo Fisher Scientific, 15520018) containing 5 mM K_3_[Fe(CN)_6_], 5 mM K_4_[Fe(CN)_6_], 150 mM NaCl, 2 mM MgCl2, 40 mM citric acid/sodium phosphate, at pH 6.0. Stained cells were washed with PBS and stored at 4 °C until imaged by light microscopy. For each sample, 500–1500 cells were analyzed.

### RNA extraction and RT-qPCR

Total RNA was isolated from cells using the RNAeasy Micro kit (Qiagen). mRNAs were reverse transcribed with the SuperScript^TM^ VI Reverse Transcriptase (Thermo Fisher Scientific, 18090050). The reaction mix consisted of 0.5mM dNTPs, 2.5 µM of Oligo d(T)20 primer, 1x SSIV buffer, 5mM of DTT, 10U/µl of Superscript^TM^ IV Reverse Transcriptase and up to 1µg template RNA. The resulting cDNAs were treated with RNAse H (Takara Bio, 2150B), to degrade possible RNA residues, and quantified by real-time RT-qPCR using the PowerUp™ SYBR™ Green Master Mix (Thermo Fisher Scientific, A25742) on a StepOne Plus RealTime PCR system (Applied Biosystems). Three independent replicates were used for each condition. Data were analysed by the StepOne Plus RT PCR software v2.1 (Thermo Fisher Scientific). TATA Binding Protein (TBP) mRNA levels were used for normalization of each target (=ΔCT). Real-time PCR CT values were analyzed using the 2^-ΔΔCt^ method to calculate the fold expression. qPCR Primers used are listed in *SI Appendix,* Table S2, including the corresponding reference.

### ATP steady-state assays

Ten thousand cells were plated on 96-well white-wall plates and were treated with 10 µM oligomycin for 1 h (for glycolytic ATP) or untreated (for total ATP levels). Cells were then tested with the CellTiter-Glo Luminescent assay (Promega), using a Tecan plate reader, according to supplier instructions.

### Transfection of expression vectors

pcDNA3.1 containing the HTRA3 ORF and a DKK tag was purchased from Genscript (OHu28769). The HTRA3 expressing vector, and empty pcDNA3.1 used as control, were transfected in MRC5-SV cells using Lipofectamine 3000 (ThermoFisher Scientific) following the manufacturer’s protocol. To select stable integrations, 48 h after transfection, 2 µg/ml puromycin was added to the culture medium, and cells were selected for more than 2 weeks. In this study, cells transfected with pcDNA3.1 are named MRC5-SV^empty^ and cells transfected with the HTRA3 pcDNA3.1 overexpressing vector MRC5-SV^HTRA3^.

### Statistical analyses

Statistical analyses were performed using the Graphpad Prism (version 6) software. The unpaired Student’s t test was applied for comparison between two groups. One-way analysis of variance (ANOVA) with post-hoc Tukey’s test was used to determine statistical significance of more than two groups. Significance indicated as p* <0.05, **<0.01, ***<0.001.

## Acknowledgements

UTechS PBI is part of the France–BioImaging infrastructure network (FBI) supported by the French National Research Agency (ANR-10–INSB–04; Investments for the Future), and acknowledges support from ANR/FBI and the Région Ile-de-France (program "Domaine d’Intérêt Majeur-Malinf") for the use of the Zeiss LSM 780 Elyra PS1 microscope. This work was supported by grants from Agence National de la Recherche, ANR, (CS_AGE, aapg2019), and ANR-DFG (Deutsche Forshungsgemeinshaft (CS_METAB, aapg2020). CFM was supported by the Pasteur-Paris University (PPU) International Ph.D. Programme, funded by the European Union’s Horizon 2020 research and innovation programme under the Marie Sklodowska-Curie grant agreement No 665807, La Ligue Contre le Cancer and Laboratoire d’Excellence (Labex) Revive.

## Author Contributions

CFM performed most experiments in this study, conceived and analysed experiments, and wrote a draft version of the paper; LC performed preliminary experiments, including mass spectrometry, and is at the origin of the hypothesis of the involvement of cathepsin B and RAC1 in POLG1 degradation; BM performed some molecular biology experiments experiments; AuS performed data acquisition and analysis of super-resolution microscopy; AlS provided patient-derived fibroblasts and discussed aspects of this work; CC performed some molecular biology experiments, conceived and analysed experiments, supervised the project, and wrote the manuscript; MR conceived and analysed experiments, supervised the project, and wrote the manuscript.

