## Supplemental Information for "HTRA3 protease-chaperone stabilizes cathepsin B for mitochondrial POLG1 depletion in human cell ageing"

**This PDF includes**

**Figures S1 to S9 and their legends, Tables S1 and S2, Supplementary References**

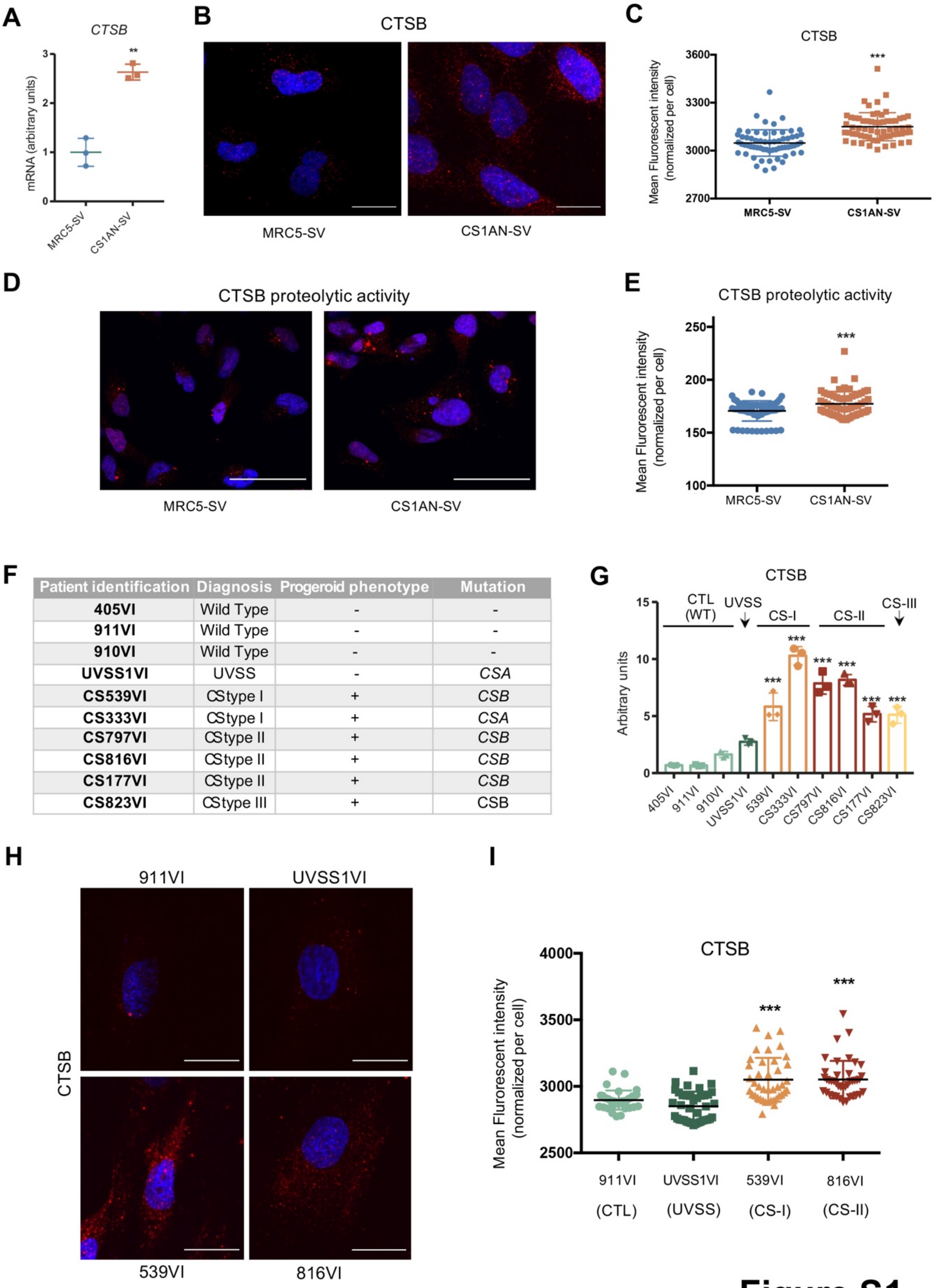


| Patient identification | Diagnosis | Progeroid phenotype | Mutation |
|---|---|---|---|
| **405VI** | Wild Type | - | - |
| **911VI** | Wild Type | - | - |
| **910VI** | Wild Type | - | - |
| **UVSS1VI** | UVSS | - | *CSA* |
| **CS539VI** | CStype I | + | *CSB* |
| **CS333VI** | CStype I | + | *CSA* |
| **CS797VI** | CStype II | + | *CSB* |
| **CS816VI** | CStype II | + | *CSB* |
| **CS177VI** | CStype II | + | *CSB* |
| **CS823VI** | CStype III | + | CSB |

**Figure S1**

**Figure S1. CTSB levels in CS primary and immortalized fibroblasts and description of WT, UVSS and CS primary skin fibroblasts.**

(**A**) RT-qPCR of *CTSB* in MRC5-SV and CS1AN-SV. (**B**) Representative confocal acquisitions of cells immunostained for CTSB (red) and counterstained with Hoechst (blue) after a z-projection into a sum slices image using ImageJ software, scale bar = 50 µM, and (**C**) quantification of the CTSB mFI (mean fluorescence intensity)/cell. Representative confocal acquisitions (**D**) and quantification (**E**) of CTSB activity (red). Cells were counterstained with Hoechst (blue) and images are the result of a z-projection into a sum slices image using ImageJ software, scale bar = 50 µM. (**F**) Characteristics of primary skin fibroblasts derived from three healthy donors (WT), one patient with UVSS and six patients with CS. (**G**) Quantification of three independent WB analysis of CTSB expression in CS patient-derived skin fibroblasts *vs.* control and UVSS. (**H**) Representative confocal acquisitions of cells in selected representative cells of the main groups of the disease immunostained for CTSB (red) and counterstained with Hoechst (blue) after a z-projection into a sum slices image using ImageJ software, scale bar = 50 µM, and (**I**) quantification of the CSB mFI/cell. RT-qPCR and WB experiments: n=3; mean ± SD. IF experiments: n=30-50 cells from 3 independent experiments; mean ± SEM. IF measurements include normalization to cell size. **$p \leq 0.01$, ***$p \leq 0.001$; based on the unpaired t test *vs.* WT MRC5-SV (panels A, C, E), or on one-way ANOVA with post-hoc Tukey's test *vs*. the mean of the controls 405VI, 911VI, 910VI (panel G) *vs*. 911VI (panel I).

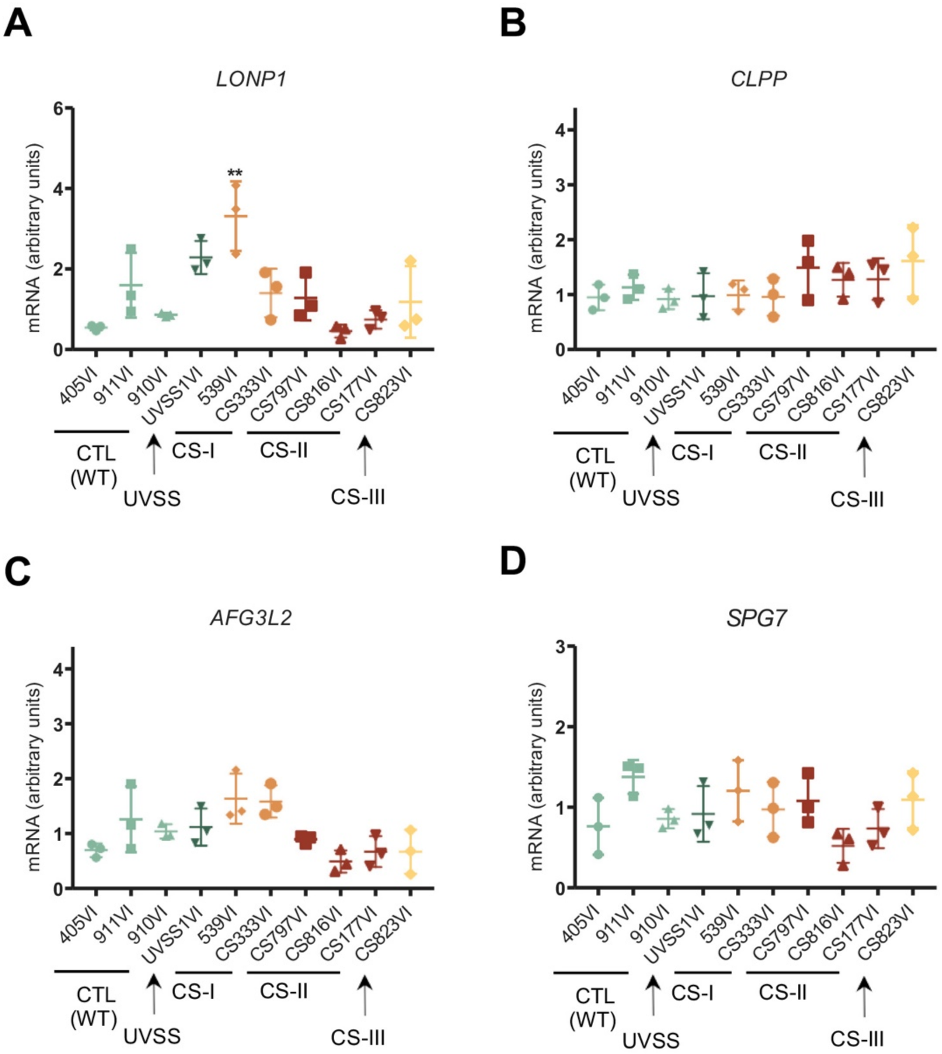


### Figure S2

**Figure S2. Expression levels of relevant mitochondrial proteases**

RT-qPCR analysis of (**A**) *LONP1* (**B**) *ClpP* (**C**) *AFG3L2* and (**D**) *SPG7* mRNA levels in WT, UVSS and CS primary fibroblasts. $^{**}p \leq 0.01$; based on one-way ANOVA with post-hoc Tukey's test *vs*. the mean of the controls 405VI, 911VI, 910VI.

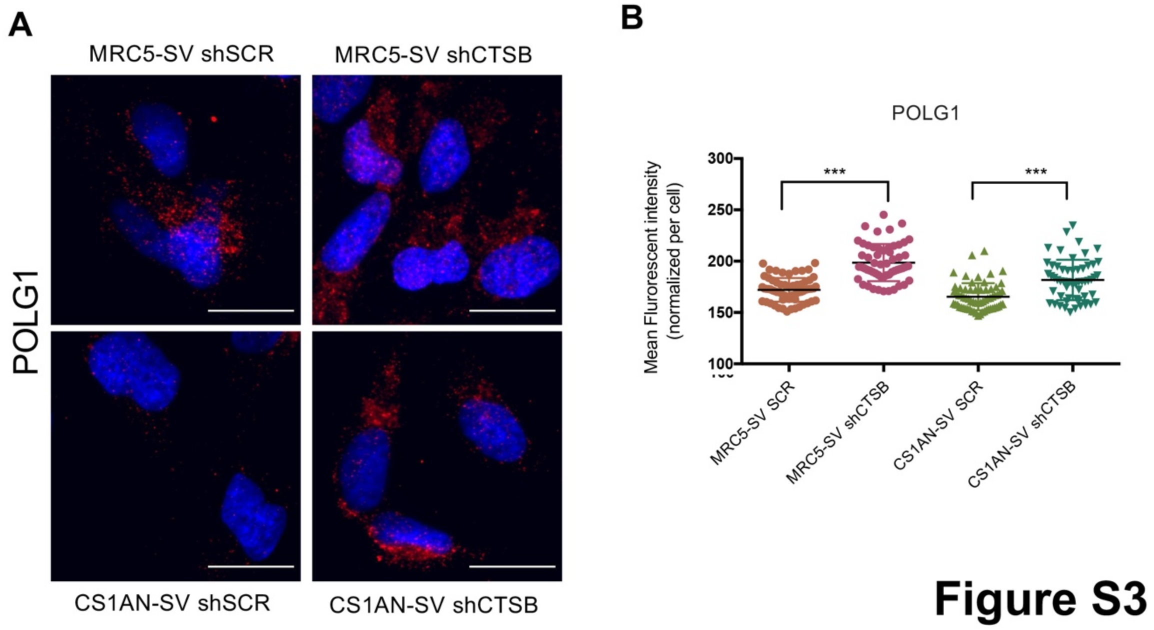


### Figure S3

**Figure S3. Immunolabeling of POLG1 and mitochondrial OXPHOS ATP levels upon CTSB downregulation**

(**A**) Representative confocal acquisitions of cells immuonstained for POLG1 (red) and counterstained with Hoeschst (blue) after z-projection into a sum slices image using ImageJ software, scale bar = 20 µM. (**B**) Quantification of the POLG1 mFI/cell. IF experiments: n=30-50 cells from 3 independent experiments; mean ± SEM. IF measurements include normalization to cell size *P ≤ 0.05, **P ≤ 0.01, ***P ≤ 0.001; based on unpaired t test comparisons to respective shSCR controls.

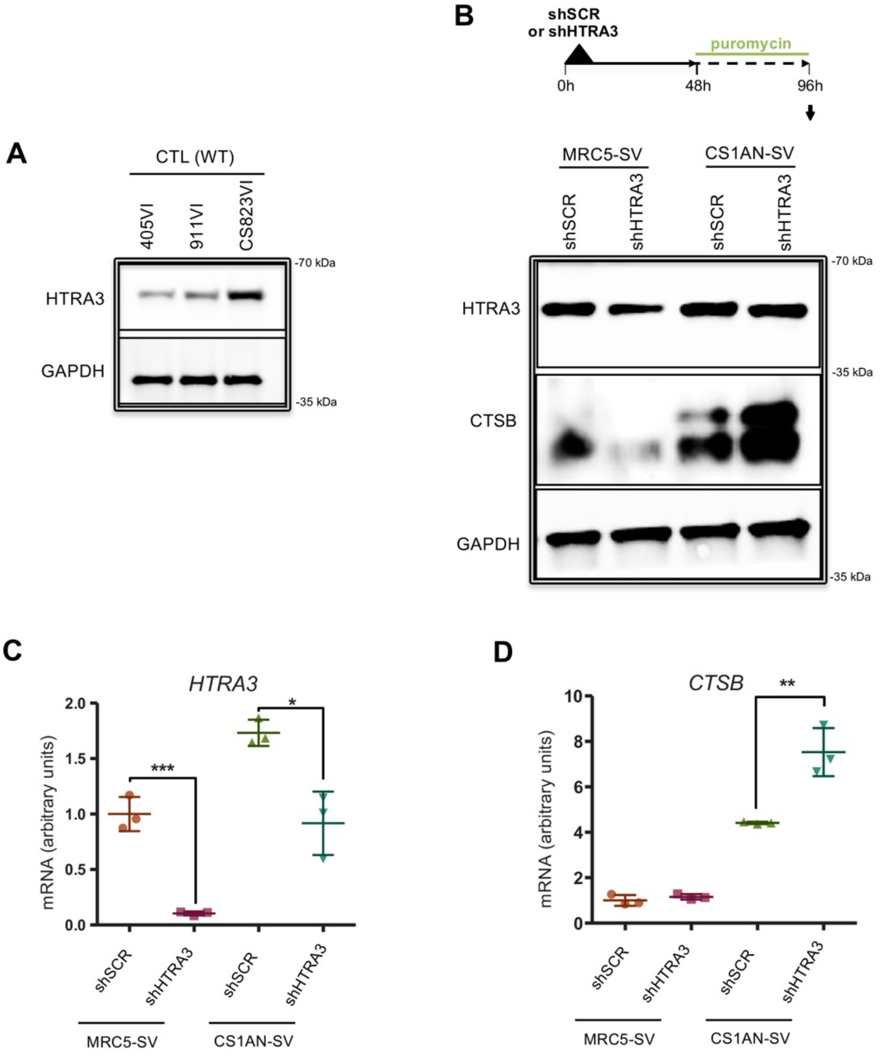


**Figure S4**

**Figure S4. Assessment of HTRA3 levels in CS-III primary fibroblasts and CTSB and RAC1 levels 96 h post-transduction of shHTRA3**

(**A**) WB analysis of HTRA3 and GAPDH (loading control) in CS-III (823) and control (405VI and 911VI) cells. HTRA3 levels in the others WT control, UVSS and CS patient-derived cells has been previously reported [1]. (**B**) Scheme showing HTRA3 silencing experiment (*upper panel panel*). WB of HTRA3 and CTSB at 96 h post shHTRA3 transduction, using GAPDH as loading control (*lower panel*). Quantitative RT-qPCR of (**C**) *HTRA3* (**D**) *CTSB* in MRC5-SV and CS1AN-SV fibroblasts knocked down for HTRA3 (shHTRA3) and scramble control (shSCR) at the same timepoint as panel b. RT-qPCR n=3 independent experiments; mean ± SD. *P ≤ 0.05, **P ≤ 0.01, ***P ≤ 0.001; based on unpaired t test comparisons to respective shSCR controls.

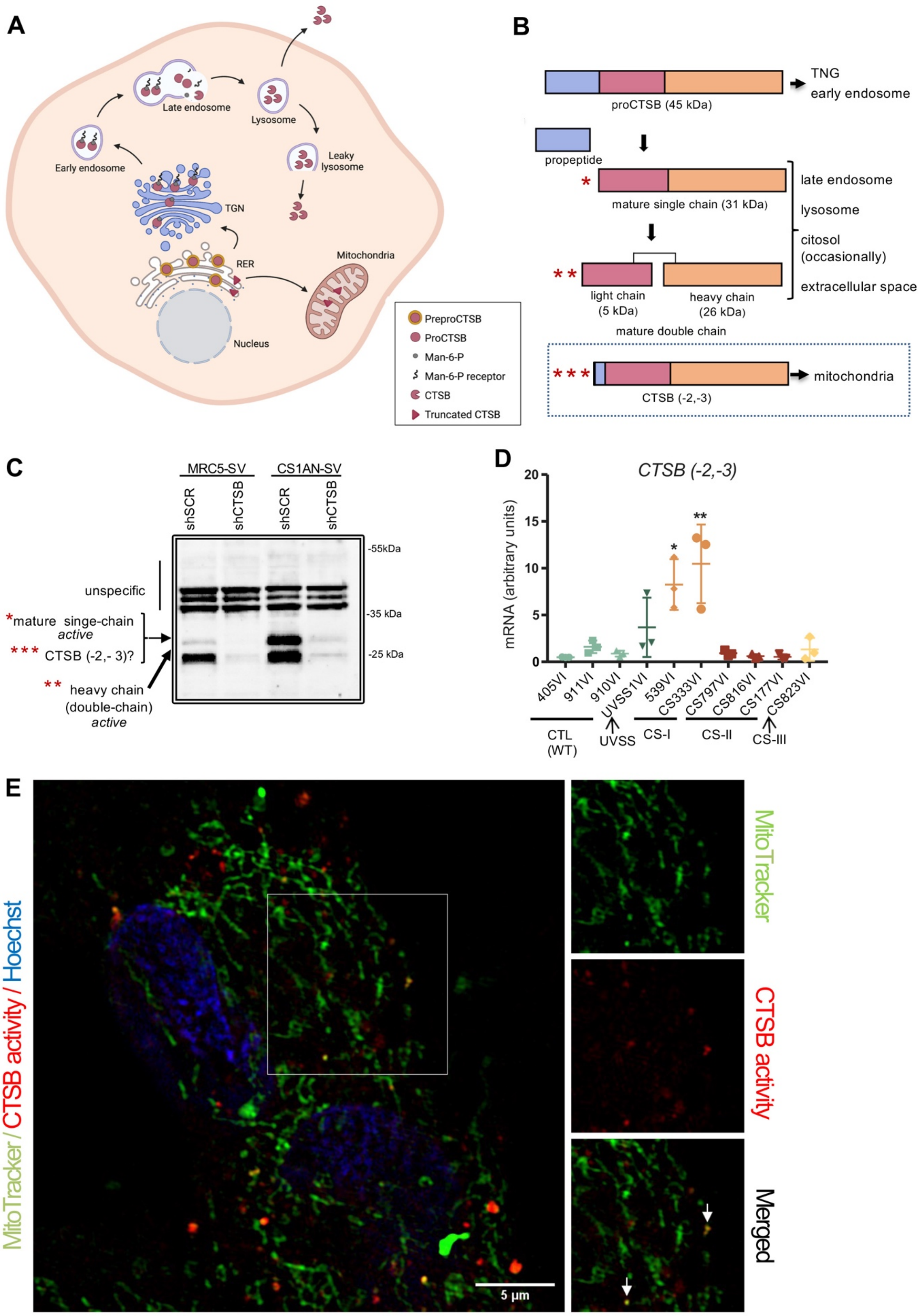
A
Late endosome
Lysosome
Early endosome
Leaky lysosome
TGN
RER
Mitochondria
Nucleus
PreproCTSB
ProCTSB
Man-6-P
Man-6-P receptor
CTSB
Truncated CTSB
B
proCTSB (45 kDa)
TNG
early endosome
propeptide
mature single chain (31 kDa)
late endosome
lysosome
citosol (occasionally)
extracellular space
light chain (5 kDa)
heavy chain (26 kDa)
mature double chain
CTSB (-2,-3)
mitochondria
C
MRC5-SV
CS1AN-SV
shSCR
shCTSB
shSCR
shCTSB
-55kDa
-35 kDa
-25 kDa
unspecific
*mature singe-chain active
*** CTSB (-2,- 3)?
** heavy chain (double-chain) active
D
CTSB (-2,-3)
mRNA (arbitrary units)
20
15
10
5
0
405VI
911VI
910VI
UVSS1VI
539VI
CS333VI
CS797VI
CS816VI
CS177VI
CS823VI
CTL (WT)
UVSS
CS-I
CS-II
CS-III
E
MitoTracker / CTSB activity / Hoechst
5 µm
MitoTracker
CTSB activity
Merged

### Figure S5

**Figure S5. Mitochondria-targeted truncated CTSB (-2, -3) expression and active CTSB**

(**A**) Schematic representation of the subcellular localization of the different forms of CTSB. (**B**) Illustration of CTSB forms. CTSB is synthesized as inactive preproenzyme (proCTSB), that is activated in the endosomal/lysosomal compartment [2]. Removal of the propeptide generates a 31 kDa mature single chain form of CTSB, which is enzymatically active. This single chain form of CTSB can be further processed by a proteolytic cleavage between residues 47 and 50, generating a heavy chain (26 kDa) and a light chain (5 kDa), that are linked by a cysteine–cysteine bond, forming a double chain form [3]. Thus, mature CTSB c exists in a single chain or a double chain form, which have been reported to have similar activities [4, 5]. CTSB (-2,-3), showed within a hatched frame, is a truncated inactive form of the enzyme that lacks the signal peptide and part of the inhibitory propeptide, and is targeted to mitochondria [6-8]. (**C**) Indication of specific CTSB forms obesrved in MRC5-SV and CS1AN-SV cells, including upon CTSB silencing (shCTSB) to show specific CTSB bands in WB. The expected migration of the mitochondrial CTSB (-2,-3) form, if present, is indicated (with a question mark). (**D**) Analysis if *CTSB (-2,-3)* mRNA levels in primary WT, UVSS, and CS patient-derived fibroblasts. *CTSB (-2,-3)* is a truncated CTSB isoform that contains a new N-terminal leader sequence that targets this protein to mitochondria instead of lysosomes. (**E**) SIM analysis of sublocalization of CTSB activity and mitochondria in MRC5-SV (one plan of a z-stack acquisition for each cell). Mitochondria were visualized using MitoTracker (green), CTSB activity using Magic Red CTSB (red), and nuclei were stained with Hoechst (blue). Scale bars = 10 µm. A 1.5x magnification is shown on the right with immunostaining for MitoTracker, CTSB activity, and merge (representative arrow for CTSB activity / MitoTracker colocalization).

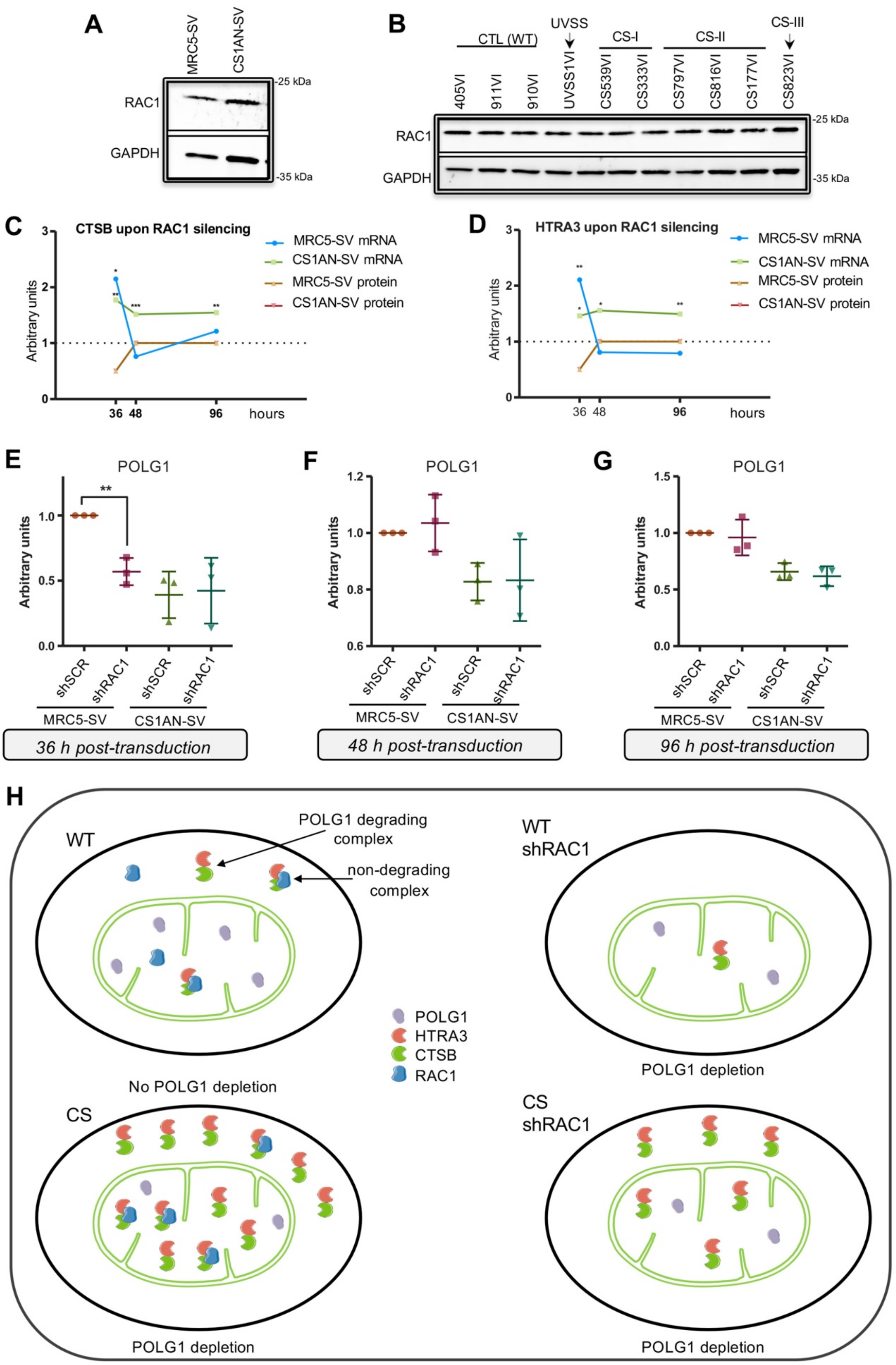

A
MRC5-SV
CS1AN-SV
RAC1
GAPDH
-25 kDa
-35 kDa
B
CTL (WT)
UVSS
CS-I
CS-II
CS-III
405VI
911VI
910VI
UVSS1VI
CS539VI
CS333VI
CS797VI
CS816VI
CS177VI
CS823VI
RAC1
GAPDH
-25 kDa
-35 kDa
C
CTSB upon RAC1 silencing
Arbitrary units
hours
MRC5-SV mRNA
CS1AN-SV mRNA
MRC5-SV protein
CS1AN-SV protein
D
HTRA3 upon RAC1 silencing
Arbitrary units
hours
MRC5-SV mRNA
CS1AN-SV mRNA
MRC5-SV protein
CS1AN-SV protein
E
POLG1
Arbitrary units
shSCR
shRAC1
MRC5-SV
CS1AN-SV
36 h post-transduction
F
POLG1
Arbitrary units
48 h post-transduction
G
POLG1
Arbitrary units
96 h post-transduction
H
WT
POLG1 degrading complex
non-degrading complex
No POLG1 depletion
WT shRAC1
POLG1 depletion
POLG1
HTRA3
CTSB
RAC1
CS
POLG1 depletion
CS shRAC1
POLG1 depletion


### Figure S6

**Figure S6. CTSB and HTRA3 protein levels are rapidly compensated upon RAC1 downregulation.**

WB analysis of RAC1 levels in (**A**) MRC5-SV and CS1AN-SV cells, and in (**B**) primary patient-derived fibroblasts; each with GAPDH as a loading control. mRNA and protein levels of (**C**) CTSB and (**D**) HTRA3 at 36 h, 48 h and 96 h after transduction with shRAC1 and shSCR in MRC5-SV and CS1AN-SV fibroblasts. mRNA levels from panels D,E,H,I,L and M in Fig. 5. Protein levels from quantification of CTSB and HTRA3 bands from panels C, G and K in Fig. 5, normalised to the respective GAPDH levels. Quantification of three independent WB analysis of POLG1 at 36 h (**E**) 48 h (**F**) and 96 (**G**) upon RAC1 downregulation. (**H**) Scheme representing RAC1-free HTRA3/CTSB digestion of POLG1 in a single mitochondrion in WT and CS cells (black ovals), in normal conditions or upon RAC1 silencing. The difference in endogenous levels of the key proteins is represented without quantitative significance. The scheme of cell composition is not at scale.

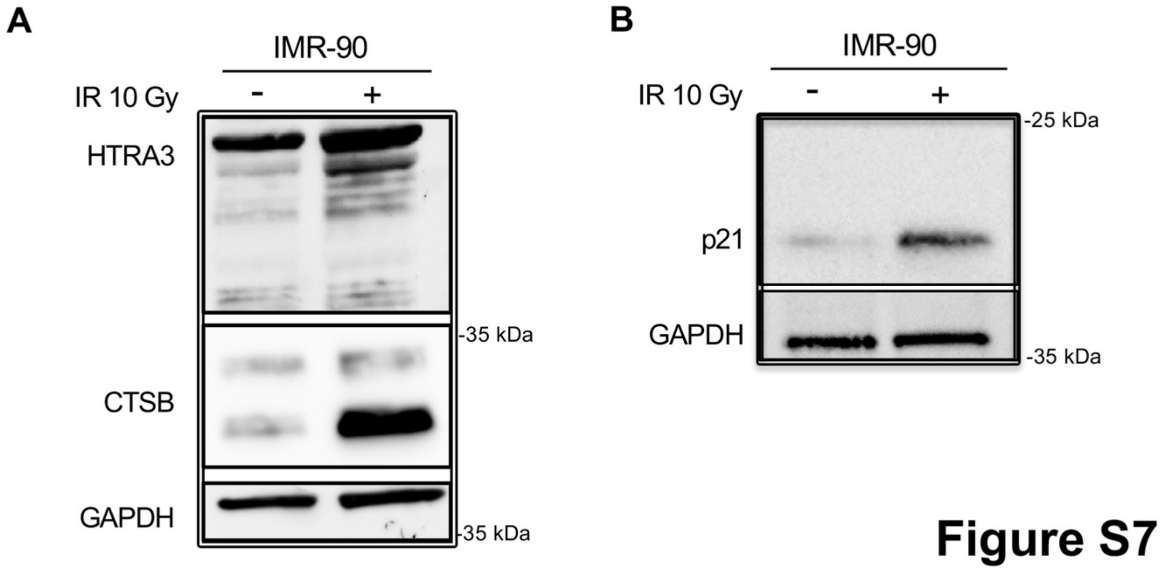


**Figure S7. CTSB overexpression recapitulated in IMR90 fibroblasts**

WB analysis of (**A**) HTRA3, CTSB, and (**B**) p21 (GAPDH staining, loading control, under each blot) in irradiated and non-IR IMR-90 fibroblasts 10 days post-irradiation.

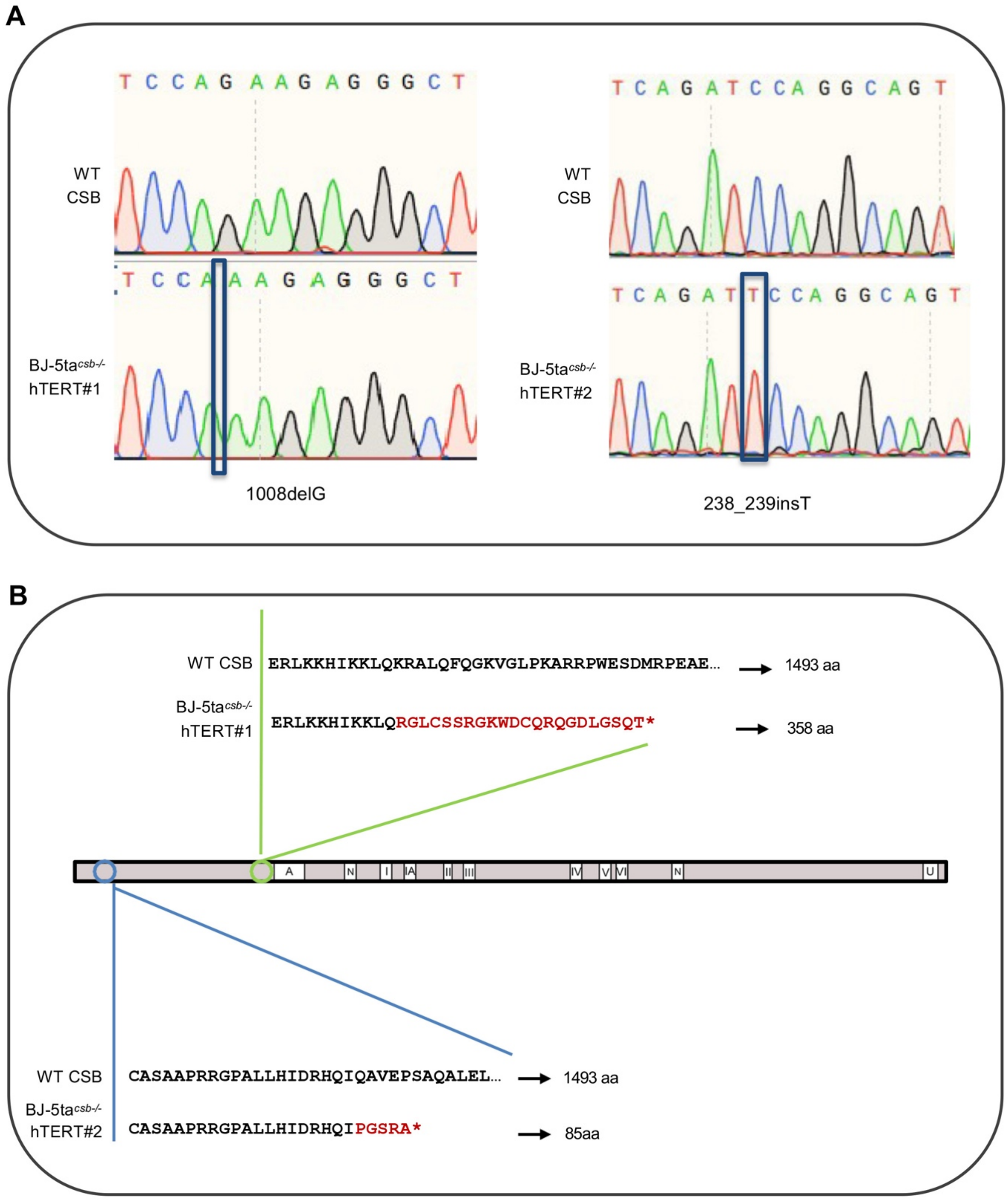
A
T C C A G A A G A G G G C T
WT CSB
T C C A A A G A G G G C T
BJ-5ta csb-/- hTERT#1
1008delG
T C A G A T C C A G G C A G T
WT CSB
T C A G A T T C C A G G C A G T
BJ-5ta csb-/- hTERT#2
238_239insT
B
WT CSB
ERLKKHIKKLQKRALQFQGKVGLPKARRPWESDMRPEAE...
1493 aa
BJ-5ta csb-/- hTERT#1
ERLKKHIKKLQRGLCSSRGKWDCQRQGDLGSQT*
358 aa
A
N
I
IA
II
III
IV
V
VI
N
U
WT CSB
CASAAPRRGPALLHIDRHQIQAVEPSAQALEL...
1493 aa
BJ-5ta csb-/- hTERT#2
CASAAPRRGPALLHIDRHQIPGSRA*
85aa

**Figure S8**

**Figure S8. Generation of isogenic BJ-5ta hTERT CSB knocked-out cells**

(**A**) Chromatograms showing the sequencing results at the target site for BJ-5ta$^{csb-/-}$hTERT#1 and BJ-5ta$^{csb-/-}$hTERT#2. Deletions and insertion sites are illustrated with squares. (**B**) Alignment CSB products resulting from mutations in BJ-5ta$^{csb-/-}$hTERT#1 (upper panel) and BJ-5ta$^{csb-/-}$hTERT#2 (lower panel) with WT protein. The size of the CSB polypeptide (aa) is indicated on the right for each case. Red *, premature stops codons; residues matching the CSB WT protein are shown in black and those resulting from frameshift mutations that do not align with the WT sequence are shown in red. CSB protein representation adapted from [9]. Domains of the CSB protein are specified: A, acidic domain; N, nuclear localisation domain; I, IA, II–VI, helicase-like domains; U, ubiquitin-binding domain.

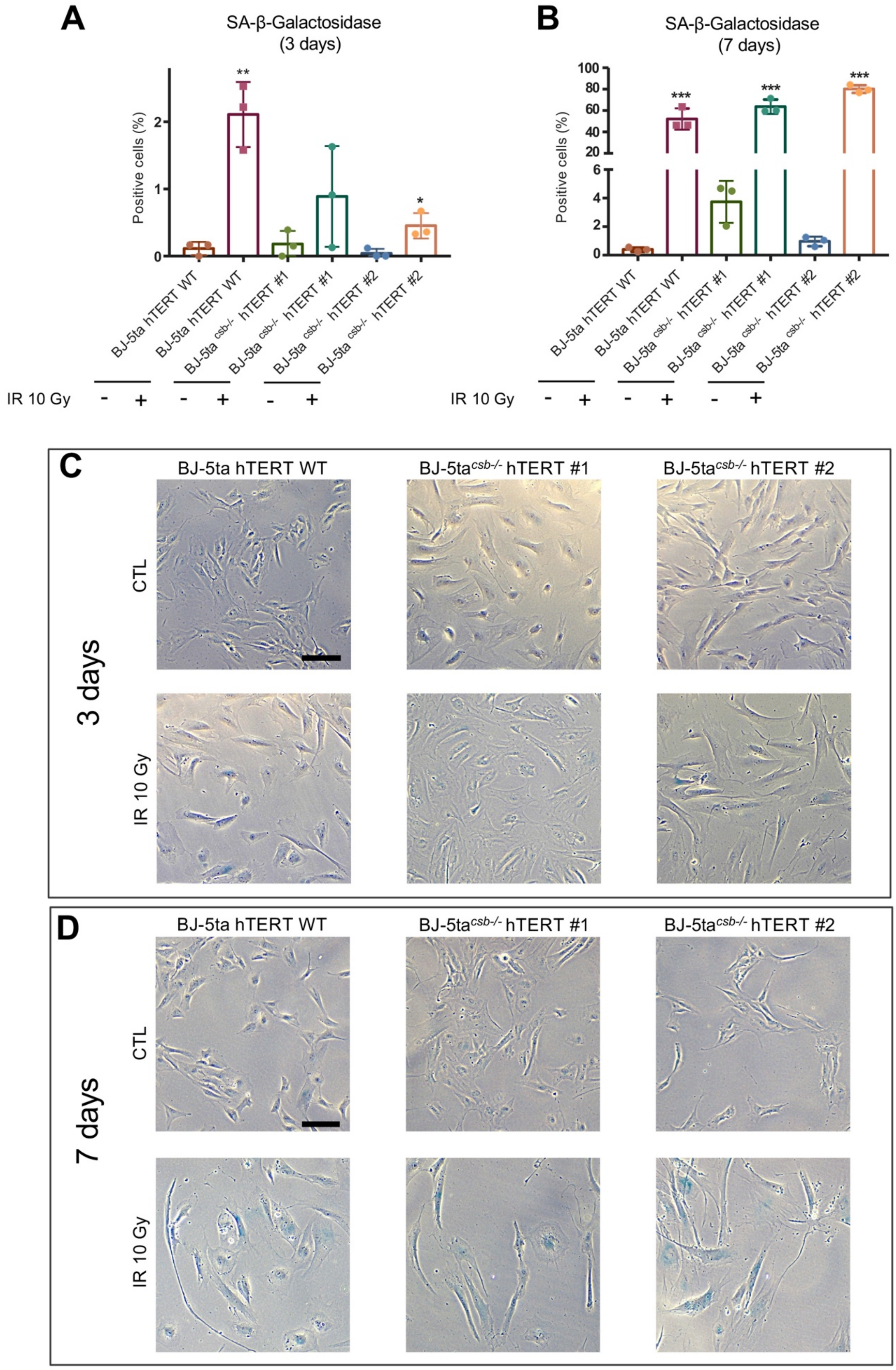
A
SA-β-Galactosidase
(3 days)
Positive cells (%)
**
*
B
SA-β-Galactosidase
(7 days)
***
BJ-5ta hTERT WT
BJ-5ta csb-/- hTERT #1
BJ-5ta csb-/- hTERT #2
IR 10 Gy
C
3 days
CTL
D
7 days

Figure S9

**Figure S9. SA-β-gal staining in CTL and irradiated BJ-5ta hTERT WT and BJ-5ta hTERT CSB knocked-out fibroblasts**

Percentage of SA-β-gal$^{+}$ cells at 3 (**A**) and 5 (**B**) days post-irradiation. *$p \leq 0.05$, **$p \leq 0.01$, ***$p \leq 0.001$; based on unpaired t test comparisons to respective non-irradiated control. Representative images of SA-β-gal+ cells at three days (**C**) and seven days (**D**) post-irradiation, scale bar = 100 μM.

#### Table S1. shRNA plasmids used for RNA interference

| Plasmid | Producer | Identifier |
|---|---|---|
| shRNA (sh Control) (SCR)-pLKO.1_Puro control | Mission shRNA Library (Sigma) | Cat# SHC002 |
| shRNA (sh CTSB)-pLKO.1_Puro CTSB | Mission shRNA Library (Sigma) | Cat# TRCN0000003658 |
| shRNA (sh HTRA3)-pLKO.1_Puro HTRA3 | Mission shRNA Library (Sigma) | Cat# TRCN0000075212 |
| shRNA (sh RAC1)-pLKO.1_Puro RAC1 | Mission shRNA Library (Sigma) | Cat# TRCN0000318432 |
| pcDNA3.1-HTRA3 | Genscript | Cat# OHu28769 |

#### Table S2. Primers used for qPCR

| Name | Sequence (5’ -3’) | Reference |
|---|---|---|
| Human CTSB Forward | CCAGGGAGCAAGACAGAGA | Giusti *et al*. [10] |
| Human CTSB Reverse | GAGACTGGCGTTCTCCAAAG | Giusti *et al*. |
| Human TBP Forward | CTCACAGGTCAAAGGTTTAC | Chatre *et al*. [11] |
| Human TBP Reverse | CGAAGCTCAACTTCCTCCC | Chatre *et al*. |
| Human HTRA3 Forward | TGGCTTCATCATGTCAGAGG | Li *et al*. [12] |
| Human HTRA3 Reverse | GGCAATGTCCGACTTCTTGT | Li *et al*. |
| Human RAC1 Forward | GCGTTGCCATTGAACTCACC | Zhou *et al*. [13] |
| Human RAC1 Reverse | GAGCTGCTACGCTCACTCCATTAC | Zhou *et al*. |
| Human p21Waf1 Forward | GAGGCCGGGATGAGTTGGGAGGAG | Yu *et al*. [14] |
| Human p21Waf1 Reverse | CAGCCGGCGTTTGGAGTGGTAGAA | Yu *et al*. |
| Human CSB Forward | CTGGAACAGGGAGTGCTTCA | Bott *et al*. [15] |
| Human CSB Reverse | ACTCCTTCTCCACGTCAACG | Bott *et al*. |
| Human CTSB(-2, -3) Forward | CAGCGCTGGGCCGGGCAC | Rose *et al*. [16] |
| Human CTSB(-2, -3) Reverse | CCAGGACTGGCACGACAGGC | Rose *et al*. |
